# Artificial intelligence reveals nuclear pore complexity

**DOI:** 10.1101/2021.10.26.465776

**Authors:** Shyamal Mosalaganti, Agnieszka Obarska-Kosinska, Marc Siggel, Beata Turonova, Christian E. Zimmerli, Katarzyna Buczak, Florian H. Schmidt, Erica Margiotta, Marie-Therese Mackmull, Wim Hagen, Gerhard Hummer, Martin Beck, Jan Kosinski

## Abstract

Nuclear pore complexes (NPCs) mediate nucleocytoplasmic transport. Their intricate 120 MDa architecture remains incompletely understood. Here, we report a near-complete structural model of the human NPC scaffold with explicit membrane and in multiple conformational states. We combined AI-based structure prediction with *in situ* and *in cellulo* cryo-electron tomography and integrative modeling. We show that linker Nups spatially organize the scaffold within and across subcomplexes to establish the higher-order structure. Microsecond-long molecular dynamics simulations suggest that the scaffold is not required to stabilize the inner and outer nuclear membrane fusion, but rather widens the central pore. Our work exemplifies how AI-based modeling can be integrated with *in situ* structural biology to understand subcellular architecture across spatial organization levels.

**One sentence summary:** An AI-based, dynamic model of the human nuclear pore complex reveals how the protein scaffold and the nuclear envelope are coupled inside cells.

## Introduction

Nuclear Pore Complexes (NPCs) are essential for the transport between the nucleus and cytoplasm, and critical for many other cellular processes in eukaryotes (*1*–*4*). Analysis of the structure and dynamics of the NPC at high resolution has been a longstanding goal towards a better molecular understanding of NPC function. The respective investigations have turned out challenging because of the sheer size, and their compositional and architectural complexity. With a molecular weight of about 120 MDa, NPCs form an extensive 120 nm-wide protein scaffold of three stacked rings: two outer rings—the cytoplasmic (CR) and the nuclear ring (NR), and the inner ring (IR). Each ring comprises eight spokes that surround a 40 to 50 nm-wide transport channel (*5, 6*). A single human NPC contains ∼30 distinct nucleoporins (Nups) in approximately 1,000 copies altogether. They arrange in multiple sub-complexes, most prominently the so-called Y-complex (*7*) arranged in head-to-tail within the outer rings (*8*). The assembly of individual sub-complexes into the higher order structure is facilitated by a yet incompletely characterized network of short linear motifs (SLiMs) embedded into flexible Nup linkers (*9*–*13*), which have been conceived as a molecular glue that stabilizes the scaffold. As a further complication, the assembled scaffold is embedded into the nuclear envelope (NE). Components of the NPC scaffold interact with the NE via amphipathic helices and transmembrane domains, and are believed to stabilize the fusion of the inner (INM) and the outer nuclear membrane (ONM) (*14, 15*). Finally, the scaffold grafts FG-Nups to the central channel, which form the permeability barrier (*16*–*18*) with their intrinsically disordered FG-rich domains, making them challenging to study using structural biology methods.

Due to these intricacies, the current structural models have severe shortcomings. In case of the human NPC, only 16 Nups, accounting for ∼35 MDa (30%) of the molecular weight of the complex are included in the models (*11, 19, 20*). Although the repertoire of atomically resolved structures of Nups has tremendously grown (*5, 6*) the respective structures often have gaps in their sequence coverage, while homology models used by many studies have intrinsic inaccuracies. For some Nups, no structures or homology models are available. Also, structural models put forward for other species were either incomplete or had a limited precision (*12, 19*–*24*). Moreover, the NPCs from many other species have a vastly reduced architectural complexity, which limits their usefulness for studying the human biology (*12, 22, 23, 25, 26*). The exact grafting sites for FG-Nups, which are crucial for understanding the transport mechanism, remain elusive. How exactly the NPC scaffold is anchored to the membrane, how it responds to mechanical cues imposed by the nuclear envelope, and if and how it contributes to shaping the membrane, remains unknown. Finally, the models remain static snapshots that do not take conformational dynamics into account.

Here, we have combined cryo-electron tomography (cryo-ET) analysis of the human NPC from isolated NEs and within intact cells with artificial intelligence (AI)-based structural prediction to infer a model of >90% of the human NPC scaffold at unprecedented precision and in multiple conformations. We demonstrate that AI-based models of Nups and their sub-complexes built using AlphaFold (*27*) and RoseTTAfold (*28*) are consistent with unreleased X-ray crystallography structures, cryo-EM maps, and complementary data. We elucidate the exact three-dimensional (3D) trajectory of linker Nups, the organization of membrane-binding domains and grafting sites of most FG-Nups in both, the constricted and dilated conformation. Our near-complete structural model enables novel types of analyses. It reveals that linker Nups perform dedicated spatial organization functions within the higher order structure. It enables molecular dynamics simulations of the scaffold embedded into explicit membrane that illustrate how the human NPC scaffold counteracts mechanical forces imposed by the nuclear membranes.

## Results

### A near-complete model of the human NPC scaffold

The completeness of the previous structural models of the human NPC was limited by the resolution of the available EM maps in both the constricted and the dilated states, and the lack of atomic structures for several Nups (*19*–*21*). To improve the resolution of the constricted state of the NPC, we subjected nuclear envelopes purified from HeLa cells to cryo-ET analysis, as described previously (*19, 21*). We collected an about five times larger dataset and applied a novel geometrically-restraint classification procedure (see Methods). These improvements resulted in EM maps with resolutions of 12 Å for the CR, 12.6 Å for IR, and 23.2 Å for NR (Fig. S1Fig. **S3**). Recent single-particle analysis of NEs isolated from *Xenopus laevis* oocytes yielded nominally higher resolved maps (*29*), but relied on spread NEs containing NPCs in only top views. Next, we obtained an *in cellulo* cryo-ET map of dilated human NPCs in the native cellular environment within intact HeLa and HEK293 cells subjected to cryo-Focused Ion Beam (cryo-FIB) specimen thinning (Fig. S1). The dilated, *in cellulo* NPC exhibits the IR diameter of 54 Å in comparison to 42 Å in the constricted state, consistent with previous work in u2os (*30*), HeLa (*31*), SupT1 (*32*), and most recently, DLD-1 cells (*20*). The quality of our *in cellulo* map is sufficient to discern the structural features known from the constricted state, such as a double head-to-tail arrangement of Y-complexes, IR subunits, and inter-ring connectors. In agreement with the previous cryo-ET maps of NPCs (*20, 22, 23, 26*), we observe increase in the distance between the adjacent spokes within the IR.

To generate a comprehensive set of structural models of human Nups, we used the recently published protein structure prediction software AlphaFold (*27*) and RoseTTAfold (*28*). We found that most of the Nups can be modeled with high confidence scores (Fig. S5). In addition, we validated the accuracy of the models by comparison to yet unreleased structures solved by X-ray crystallography for human Nup358, Nup93, Nup88, Nup98, and of Nup205 and Nup188 from *Chaetomium thermophilum* (Fig. S6). The X-ray structures were provided to us by the Hoelz laboratory (*33, 34*) and were not used as input for the modeling procedure. The AI-based models also excellently fit our EM densities (Fig. S7A-E). With full-length models at hand, we could identify the positions of Nup205 and Nup188 within the scaffold, which could not be unambiguously located in the previous hNPC cryo-ET maps (see below). The AI-predicted conformation of the N-terminal domain of Nup358 is more consistent with the observed electron density as compared to the two X-ray structures (Fig. S7H). The Nup358 localization is in agreement with previous analysis (*21*) and the secondary structure in the *Xenopus* EM map (*29*) (Fig. S8). The full-length model of ELYS, for which thus far only the N-terminal ? -propeller could be placed (*21*), fits the EM map as a rigid body (Fig. S7E) and confirms its binding site to each of the Y-complexes in the NR. The models of Nups in the CR agree with the secondary structure observed in the *Xenopus* cryo-EM map (Fig. S8).

The capacity of AI-based structure prediction tools to identify and model protein interfaces with high accuracy has recently been demonstrated (*35, 36*). We therefore attempted to model Nup interfaces using the ColabFold software, a version of AlphaFold adopted for modeling protein complexes (*36*). We found that ColabFold predicts several Nup sub-complexes with inter-domain confidence scores that correlate with the accuracy of the models (Fig. S9-Fig. **S11**). The models of these sub-complexes did not only reproduce the respective, already available X-ray structures X-ray structures, but also agreed with the yet unreleased X-ray structures (*33, 34*). Specifically, X-ray structures of *C. thermophilum* Nup205 and Nup188 in complex with Nup93 as well as Nup93 in complex with Nup35 are consistent with the human ColabFold model (Fig. S6). These structures represent proteins in complex with the respective SLiMs and form relatively small interfaces. However, also for larger sub-complexes we obtained structural models that convincingly fit our cryo-ET maps (Fig. S11). For example, the structure predicted for the so-called central hub of the Y-complex was consistent with the organization seen in fungal X-ray structures and explained additional density within the cryo-ET map specific to the human NPC. The new model of the Y-complex hub includes a previously unknown interaction between Nup96 and Nup160 (Fig. S9). ColabFold predicted the structure of the Nup62 complex with excellent accuracy (Fig. S11) even though no structural templates were used for modeling. We were also able to obtain a trimeric model of the small arm of the Y-complex comprising Nup85, Seh1, and Nup43. The model fits the EM map very well, confirming the known structure of Nup85-Seh1 interaction, and revealing how Nup43 interacts with Nup85 (Fig. S11). In the case of the Nup214 complex, for which no structures are available, the ColabFold model is highly consistent with the rather uniquely shaped EM density (Fig. S11). A previously unknown interface between Nup214 and Nup88 that is biochemically validated in (*33*) has also been predicted correctly (Fig. S6).

With the cryo-EM maps and the repertoire of structural models of individual Nups and their sub-complexes, we built a nearly complete model of the human NPC scaffold (Fig. 1A). We used the previous model (*19, 21*) as a reference for modeling the scaffold of the constricted state and replaced all previously fitted domains with human AlphaFold and ColabFold models. We then added the remaining newly modeled subunits by systematic fitting to the EM map and refinement using the Assembline program (*37*). In addition to fitting the models, we have added several disordered linkers that connect spatially separated domains and SLiMs within the NPC. We then built the model of the dilated state by fitting the constricted NPC map into the dilated NPC map and refining the fits using Assembline. The resulting models (Fig. 1) include 25 out of ∼30 Nups, and account for 70 MDa of the molecular weight of the NPC (>90% of the scaffold molecular weight) in comparison to 16 Nups and 35 MDa (46% of the scaffold) of the previous model, and largely account for the EM density observed in the constricted and dilated states.

**Fig. 1.**
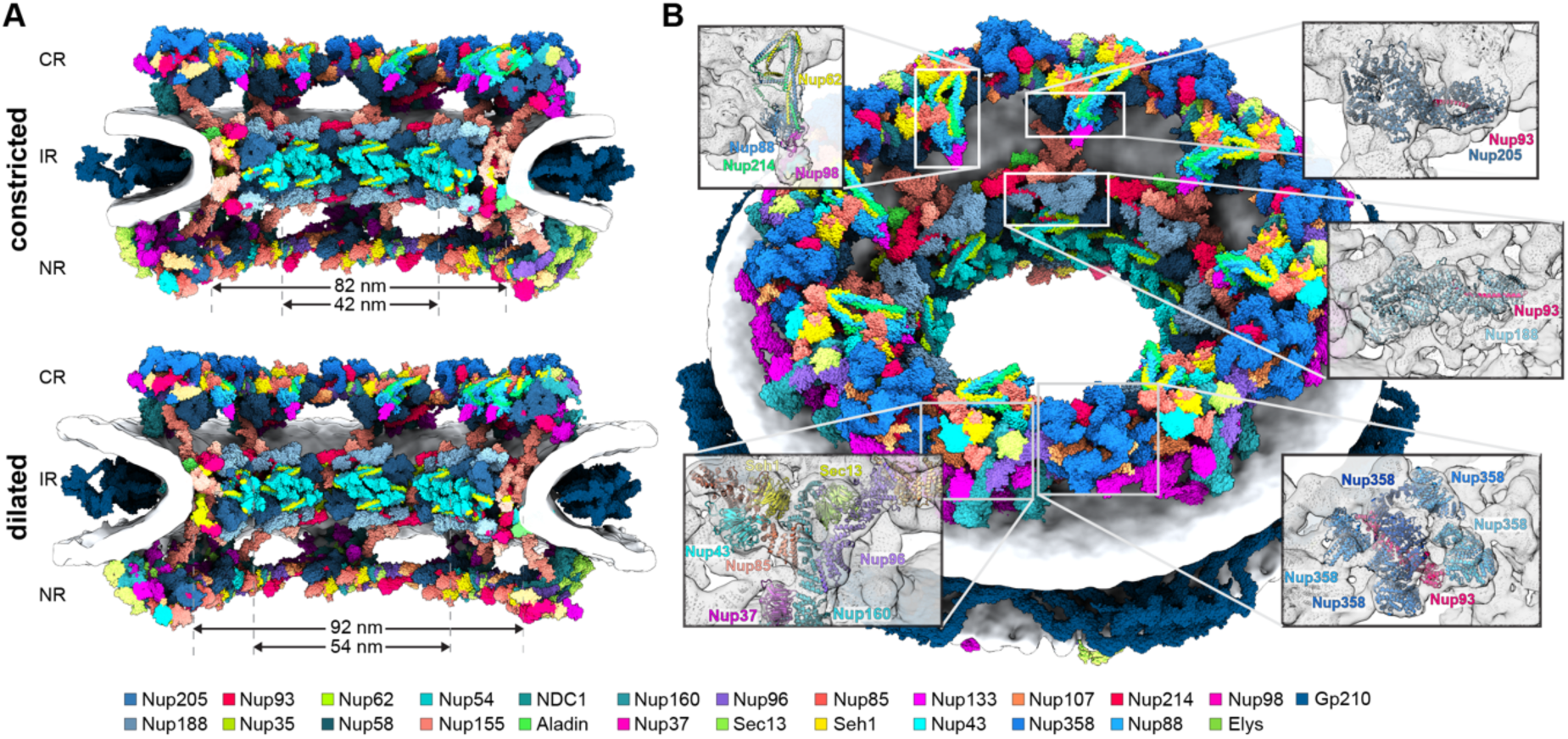
Scaffold architecture of the human NPC. **(A)** The near-complete model of the human NPC scaffold is shown for the constricted and dilated states as cut-away views. High-resolution models are color-coded as indicated in the color bar. The nuclear envelope is shown as a gray isosurface. **(B)** Same as **(A)** but shown from the cytoplasmic side for the constricted NPC. The insets show individual features of the CR and IR enlarged with secondary structures displayed as cartoons and superimposed with the isosurface-rendered cryo-ET map of the human NPC (gray).

The near-complete scaffold model yields novel insights into the organization of the human NPC (Fig. 1). It resolves previous ambiguities (*11, 19, 21, 29*) in localization of similarly shaped Nup205 and Nup188. Within the IR, Nup188 localizes to the outer, while Nup205 localizes to the inner sub-complexes, consistent with previous analysis in other species (*12, 22, 23*). We furthermore localized two copies of Nup205 in the CR and one in the NR (*29*), thus resolving previous ambiguities (*11, 19, 21*). Two previously undetected copies of Nup93 bridge the inner and outer Y-complexes both in the CR and NR, with an inherent C2 symmetry. This observation is consistent with biochemical experiments that initially identified interactors of Nup93 in the outer rings (*38*). The copy of Nup93 in the CR is located underneath the Nup358 complex, further corroborating a role of Nup358 in stabilizing the higher order structure (*21*). Yet another copy of Nup93 that is unique to the CR bridges the inner Y-complexes from two consecutive spokes. This is consistent with an additional copy of Nup205 in the CR as compared to the NR, since Nup93 and Nup205 heterodimerize through a SLiM within the extended N-terminus of Nup93 (this study and in (*33, 34*)). The AI-based model of the Nup214 sub-complex interacts with Nup85 pointing towards the central channel, likely to optimally position the associated helicase crucial for mRNA export.

### Linker Nups fulfill dedicated roles of spatial organization within the higher-order assembly

Since the exact 3D trajectory of the linkers through the NPC scaffold was unknown, it remained difficult to understand their precise structural role beyond conceptualization as molecular glue. In our model, AI-based models of human Nup-SLiM subcomplexes allowed us to map the anchor points of the linkers to the scaffold. The AI-based models correctly recapitulated SLiM interactions known from X-ray structures but also revealed previously unknown additional human Nup-SLiM interactions. In comparison to the X-ray structures, the AI-based models more extensively covered the structured domains, thus reducing the length of the linkers connecting between the anchor points and restricting their possible conformational freedom within our model. We therefore traced the linkers as ensembles of their approximate paths in explicit atomic representation using coarse-grained modeling followed by atomic refinement (Supplementary Materials and Methods).

The resulting connectivity map (Fig. 2) reveals that the Nup35 linker regions bridge between neighboring spokes of the IR. In our model, the Nup35 dimer is positioned into previously unassigned EM density between spokes, and each of the two copies reaches out with its SLiMs to Nup155 and Nup93 of the adjacent spokes (Fig. 2A). The amphipathic helices of Nup35 and a SLiM binding to Aladin (see below) limit the distance to the membrane. A Nup35 bridge between adjacent spokes is the only arrangement geometrically possible for both the constricted and the dilated NPC state. These bridges are consistent with the previous interaction and modeling studies in fungi (*10, 12, 24*) and the biochemical analysis of the human proteins (*34*). It strongly suggests that the Nup35 dimer, which is critical during early NPC biogenesis (*39*), functions as an architectural organizer for the IR membrane coat in a horizontal direction along the membrane plane.

**Fig. 2.**
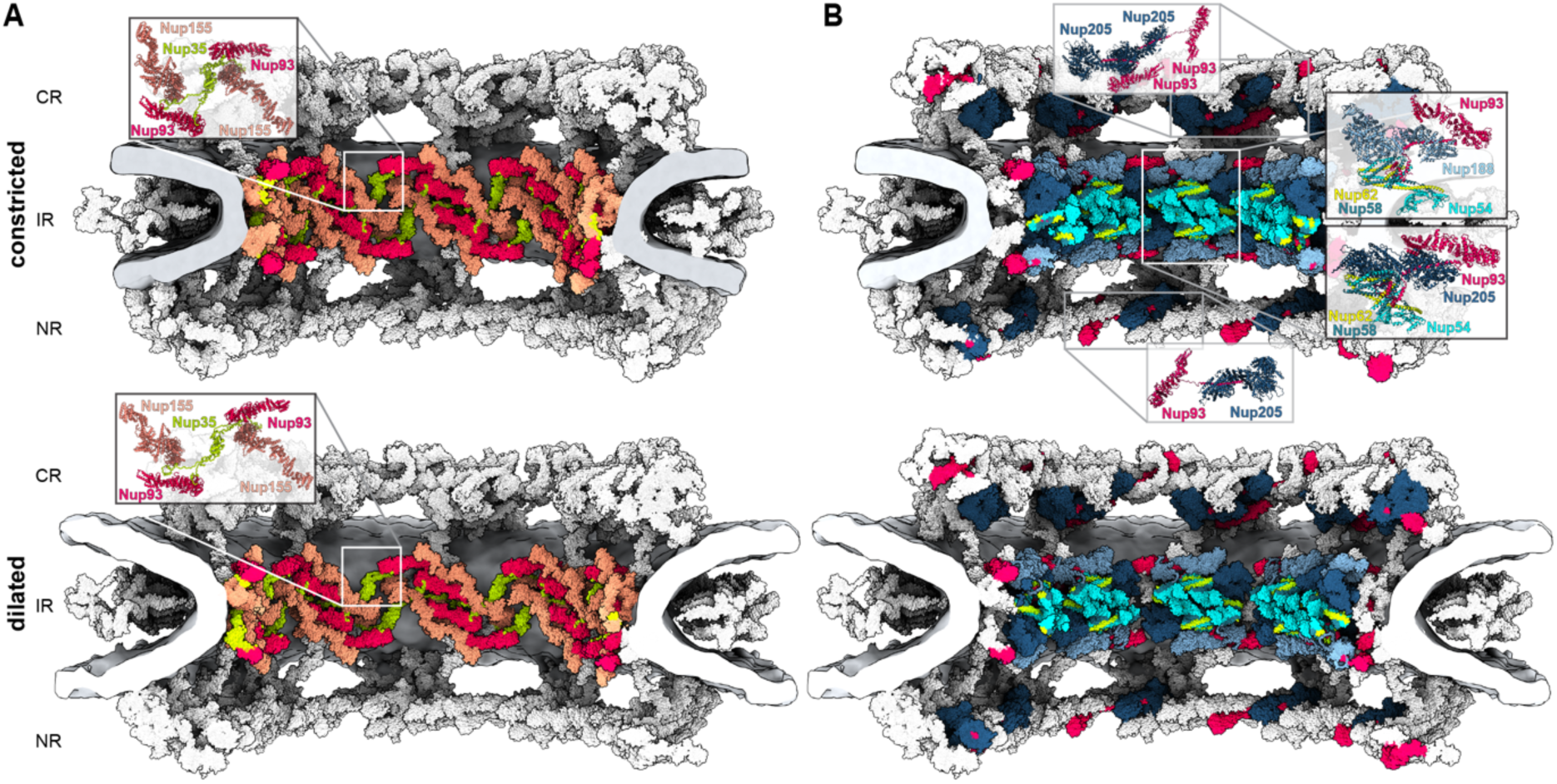
The connectivity of protein linkers within the human NPC. **(A)** The Nup35 dimer interconnects adjacent spokes across different sub**-**complex species, thus facilitating cylindrical assembly of the IR in both constricted (top) and dilated (bottom) states. The Nup205, Nup188, Nup62 complex, and the N-terminus (aa. 1-170) of Nup93 are hidden from view to expose Nup35. **(B)** The N-terminus of Nup93 isostoichiometrically connects the subunits Nup205, Nup188, Nup62, Nup58 and Nup54 within the same sub-complex species. Insets show Nup93 connectivity, highlighting its interaction with two copies of Nup205 in the CR, two copies each with Nup205 and 188 in the IR and a single copy of Nup205 in the NR. The respective subunits are shown color-coded as in Fig. 1, while all other subunits and the nuclear membranes are shown in gray.

In contrast, the connectivity map demonstrates that the linkers at the N-terminus of Nup93 copies that connect anchor points at Nup205 or Nup188, and Nup62 complex in the IR, cannot reach across spokes in either the constricted or the dilated state, and thus connect subunits within a single IR sub-complex inside the same spoke. This is the only geometrically possible solution we found for all four instances of Nup93 in the IR, with the two outer copies binding to Nup188 and the two inner copies binding to Nup205. Thus, Nup93 acts as an architectural organizer within, but not across spokes. In the CR and NR, the linkage between Nup93 and Nup205 is geometrically possible, emphasizing the similar architectural design of the respective complexes. Thereby, the Nup93 SLiM that binds the homologs Nup62 complex could also facilitate linkage between the Nup93 and the Nup214 complex, although the corresponding structural information is still missing. The duplication of Nup205 and Nup93 in the CR is suggestive of yet another copy of the Nup214 complex that is not well resolved in the EM map of the constricted state, and thus remains to be further investigated.

In conclusion, the individual linker Nups specialize in dedicated spatial organization functions responsible for distinct aspects of assembly and maintenance of the NPC scaffold architecture.

### A transmembrane interaction hub organizes the interface between outer and inner rings

Several types of structural motifs associate the NPC scaffold with the membrane. The spatial distribution of the amphipathic helices and membrane-binding loops harbored by Nups 160, 155, and 133 across the scaffold has been previously revealed (*19*). The analysis of the protein linkers in our new model allowed the mapping of an approximate location of the amphipathic helices of Nup35. In addition to these motifs, the human NPC further contains three transmembrane Nups, the precise location of which remained unknown.

Among the three human transmembrane proteins, NDC1 is the only one that is conserved across eukaryotes (*40*). NDC1 is known to interact with the poorly characterized scaffold Nup Aladin (*41, 42*). We confirmed this interaction using proximity labeling mass spectrometry with BirA-tagged Aladin and identified NDC1 and Nup35 as the most prominently enriched interactors (Fig. S12). NDC1 is predicted to comprise six transmembrane helices followed by a cytosolic domain containing mainly α-helices, while Aladin is predicted to have a ? -propeller fold. Structures of both NDC1 and Aladin, however, remain unknown. Using AlphaFold/Colabfold, we could model the structures both as monomers and a heterodimeric complex with high-confidence scores (Fig. S5, Fig. S9). Systematic fitting of the hetero-dimeric models to the EM map unambiguously identified two locations within the IR (Fig. S11). The EM density was not used as a restraint for modeling but matches the structure of the model very well and is consistent with the only patches of density spanning the bilayer (Fig. 3, Fig. S11), therefore further validating the model. The two locations are C2-symmetrically equivalent across the nuclear envelope plane, thus assigning two copies of Aladin and NDC1 per spoke, corroborating experimentally determined stoichiometry (*43*). The identified locations are close to the membrane-binding N-terminal domains of Nup155 and amphipathic helices of Nup35, which is also consistent with our proximity labeling data (Fig. S11) and previous functional analysis (*44*). The proximity labeling data also identified Gle1, which was previously shown to interact with Nup155 (*45*). Using AlphaFold we predicted an interaction between the C-terminus of Nup155 and the N-terminal SLiM in Gle1 with high confidence scores (Fig. S12B). We also predicted an Aladin-binding SLiM in Nup35, located between the amphipathic helix and Nup155-binding SLiM (Fig. 3, Fig. S9). We therefore propose that together with Nup155, Nup35, Aladin and NDC1 form a transmembrane interaction hub that anchors the inner membrane coat of the IR and orients the Nup155 connectors towards the outer rings. The central position of Aladin within the NPC might explain functional consequences of mutations in the *Aladin*/*AAAS* that are implicated in the AAA syndrome (*46*– *48*) and is consistent with the absence of Aladin in fungi, which lack the Nup155 connectors (*25*).

**Fig. 3.**
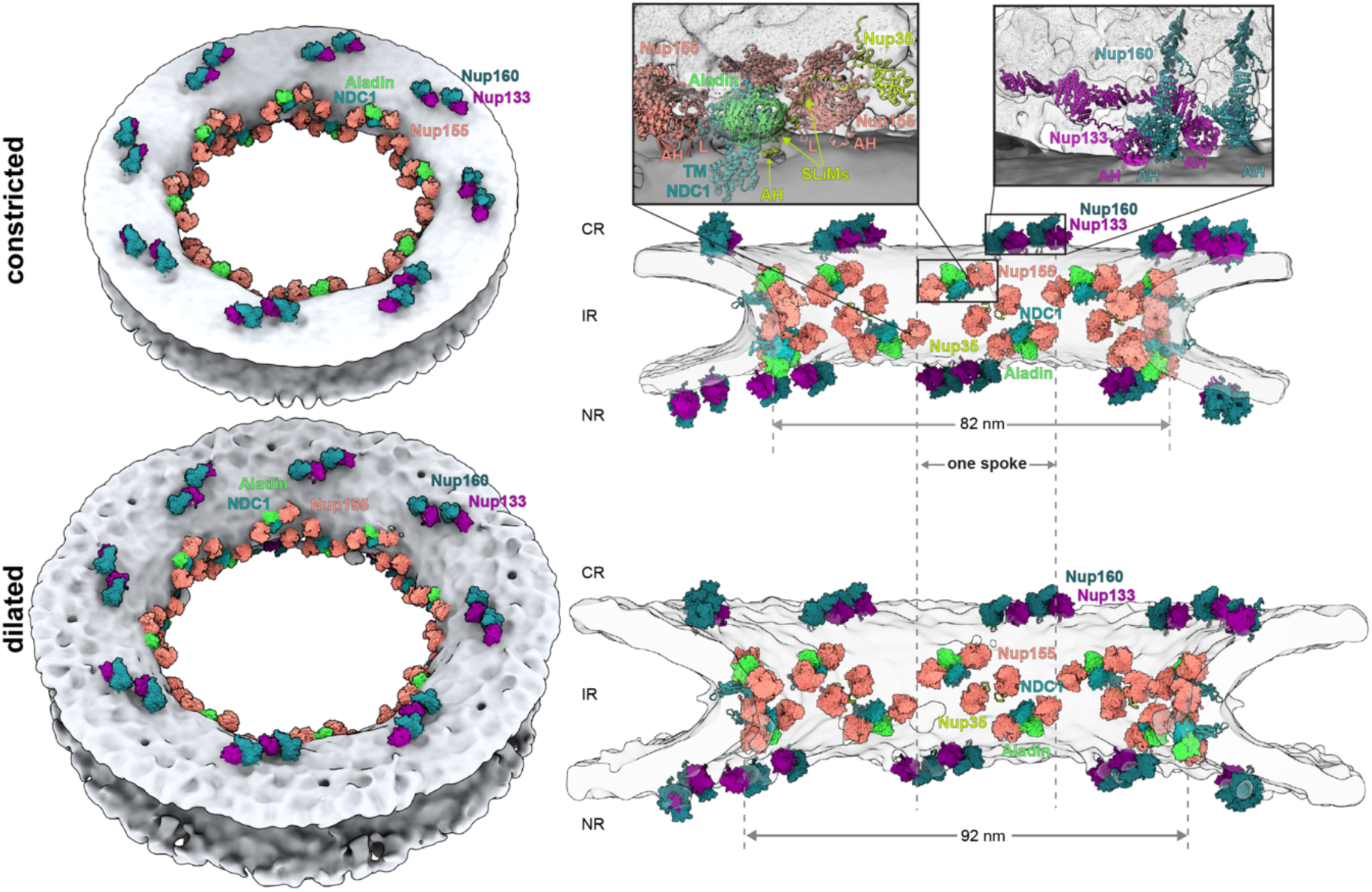
The membrane-anchoring motifs of the human NPC are distributed over the entire scaffold. The membrane-binding b-propellers of the Y-complex and IR complex are shown color-coded and arranged as pairs of the respective inner and outer copies. Aladin and NDC1 form a transmembrane interaction hub with the inner and connector copies of Nup155, which is shown enlarged in the inset in the cut-away side view (right). The nuclear membranes are shown as a gray isosurface.

We next examined the structure of Gp210, which contains a single-pass transmembrane helix and is the only Nup that primarily resides in the NE lumen. This Nup is composed of multiple immunoglobin-like domains and is thought to form a ring around the NPC within the NE lumen (*49*), but the structure of the ring has only been modeled in fungi. We used RoseTTAfold to model full-length Gp210 and obtained an elongated model with clearly defined interfaces between consecutive domains. Already as a rigid body, this model fitted the density sufficiently well to allow tracing Gp210 monomers in the cryo-EM map of the *Xenopus laevis* NPC, which is superbly resolved in the luminal region (*50*). Thus, we could assign eight copies of Gp210 per spoke (Fig. S13). Modeling of individual Gp210 fragments and inter-Gp210 interactions with AlphaFold/ColabFold (Fig. S13) led to an overall model that explained the entire density of the luminal ring of the Xenopus cryo-EM map, including the C-terminal transmembrane helix. This helix is long enough to span the NE and reaches the IR in the vicinity of the NDC1/Aladin/Nup155/Nup35 transmembrane interaction hub. This is in good agreement with known interactions of Gp210 homologs (*51*) and our proximity labeling data (Fig. S12). The model of the Gp210 ring also matches the luminal density visible in our *in cellulo* EM map, allowing us to model the ring in the context of both constricted and dilated NPC (Fig. 1, Fig. S13). To further confirm the Gp210 assignment in human cells, we deleted *gp210* in HEK293 cells using CRISPR/Cas9 and analyzed the structure of the NPCs *in cellulo* using cryo-ET. The resulting map indeed showed a lack of the luminal ring density (Fig. S4). The NPC scaffold in the resulting map appears overall unchanged, including its diameter, suggesting the Gp210 is not required for faithful NPC assembly.

Taken together, our model includes all known membrane-binding domains except for the cell-type specifically expressed Pom121, the precise location of which within the NPC remains unknown, and neither AlphaFold or RoseTTAfold could build structural models with high confidence. The resulting membrane association map reveals that the membrane-binding ? - propellers of the Y-complex (Nup160 and Nup133) and the IR (Nup155) are distributed as multiple pairs over the entire scaffold, whereby they follow a well-defined pattern. They form an overall Z-shaped outline within an individual spoke (Fig. 3). The NDC1/Aladin/Nup155/Nup35 membrane-binding hub is situated at the interface of the IR with the outer rings and is distinct from the additional Nup155 pair at the NE symmetry plane. The membrane-binding motifs arrange in similar clusters in both the constricted and dilated state. Their relative arrangement is not changing uniformly during dilation, rather the relative distances within the spokes remain constant while the spacing of the spokes increases (Fig. 3).

### The NPC scaffold prevents membrane constriction in the absence of membrane tension

It has been a longstanding view that the scaffold architecture of the NPC has evolved distinct membrane-binding motifs to stabilize the membrane at the fusion of the INM and ONM (*52, 53*). To test the contribution of the scaffold architecture to membrane curvature, we used molecular dynamics (MD) simulations using a coarse-grained Martini force field (*54, 55*). We first simulated a double-membrane pore without proteins with an initial pore diameter and membrane spacing as seen in the constricted NPC cryo-ET map. We found that the pore constricts during 1 μs simulations and stabilizes once the radii of the INM/ONM and the NE hole are the same, such that the mean curvature nearly vanishes (Fig. 4A, Fig. S15, Video S1). This is in line with Helfrich membrane elastic theory, which predicts a catenoid-like pore shape with equal radii of curvature at the pore center as the lowest energy structure, and an energetic cost of about 200 *kBT* to widen the pore (Supplementary Notes). Interestingly, the opening of the relaxed double-membrane pore is considerably smaller than even the most constricted NPC conformations. The NPC scaffold thus keeps the pore wider than it would be without the scaffold.

**Fig. 4.**
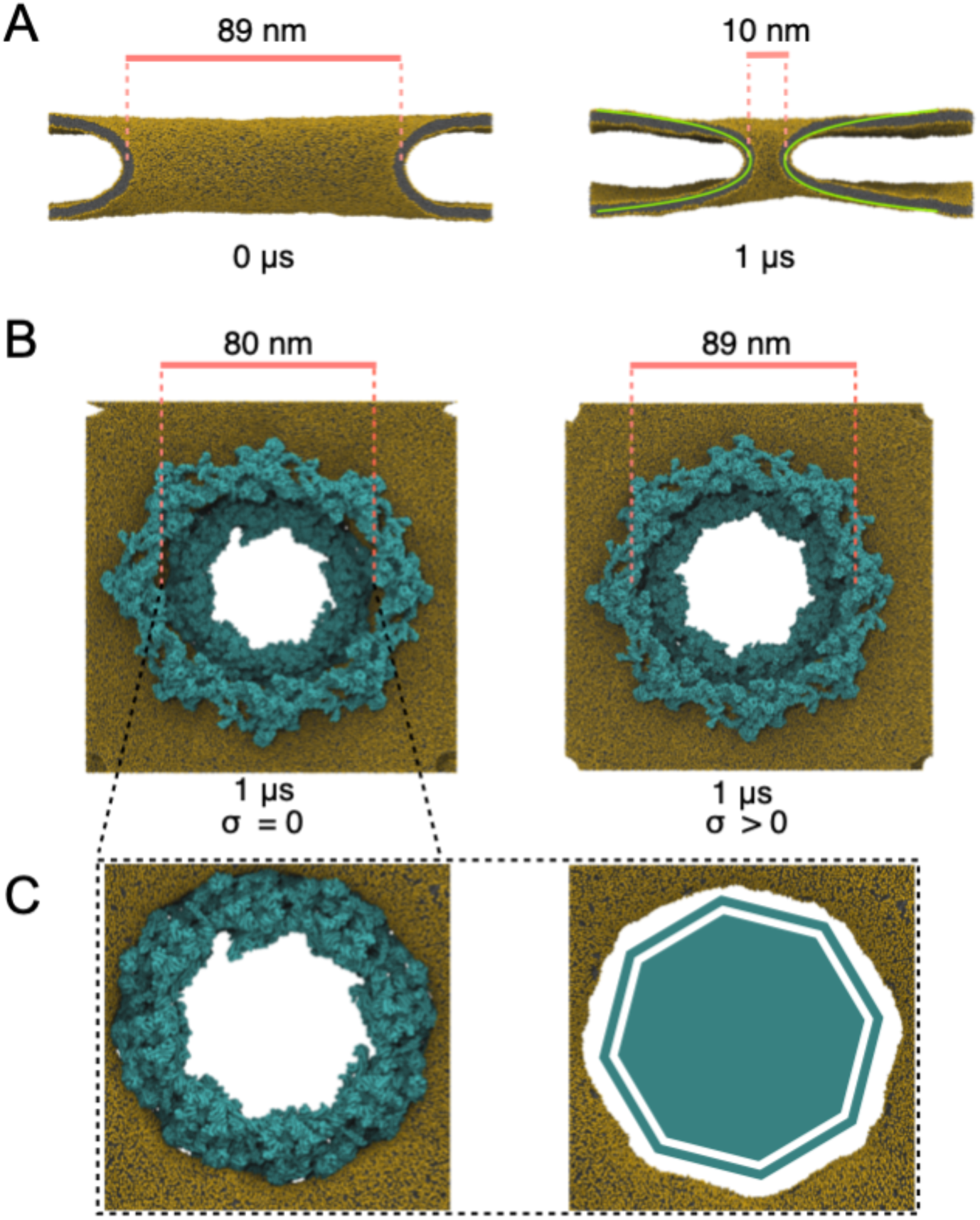
Dynamics of the NPC from molecular simulations. **(A)** An isolated half-toroidal double-membrane pore shaped initially as in the tomographic structure of the constricted NPC (left) tightens over the course of 1 μs of MD (right) toward the catenoid-like shape (green) predicted by membrane elastic theory. Shown are cuts along the axis of the double-membrane pore with lipid headgroups and tails in gold and gray, respectively. The solvent is not shown. (B) The NPC (cyan) widens by ∼10% in response to lateral membrane tension (right; ΔP=2 bar) compared to a zero-tension simulation (left; ΔP=0). Shown are snapshots of the relaxed structures after 1 μs of MD. (C) The membrane fits tightly around the NPC inner ring (cyan, left; ΔP=0) and forms an octagonally-shaped pore (right, NPC not shown).

These findings would predict that the nuclear membranes push against the NPC scaffold even in the most constricted state, which is in agreement with experimental data (*23*). To examine the effects of this tension on the NPC, we generated an NPC scaffold model with explicit membrane and water as a solvent, and ran 1 μs MD simulations (Fig. 4B, Fig. S15-Fig. **S19**, Videos S2-3). In these simulations, we found that the membrane pore wrapped tightly around the IR plane, adopting an octagonal shape (Fig. 4C). Similarly tight wrapping and octagonal shapes have been seen in the previous EM analyses of NPCs (*21, 56, 57*). We also observed that the diameter of the NPC scaffold constricted by about 9% (Fig. 4B). We attribute this tightening primarily to mechanical tension in the pore widened beyond the catenoid shape. First, we observed similar contraction in simulations with rescaled protein-protein interactions (Fig. S19). Second, by applying lateral tension on the double membrane, we could maintain the pore width or widen it (Fig. 4B, Fig. S15B). At even higher tension, the membrane spontaneously detached from the NPC scaffold (Fig. S20). Taken together, our data support a model in which the role of the NPC scaffold is not to stabilize the membrane fusion *per se*, but rather to widen the diameter of the membrane hole without necessitating a wider envelope.

## Discussion and conclusions

We have built a near-complete model of the human NPC scaffold in the constricted state (smaller diameter) as adopted in purified nuclear envelopes and in the dilated state as adopted in cells, whereby recent work in fungi has identified constricted NPCs inside of cells under specific physiological conditions (*23*). Our model includes multiple previously unassigned domains and proteins, resolves long-standing ambiguities in alternative Nup assignments, lays out a connectivity map of the protein linkers across the NPC scaffold, maps out the membrane-anchoring motifs, and provides a high-quality basis for further investigations of NPC dynamics and function. Our analysis demonstrates that coarse-grained dynamic model is of sufficient quality for molecular simulations, which in future could quantitively and predictively describe how the NPC interplays with the nuclear membrane and how it responds to mechanical challenges. The model also provides a more accurate starting point for simulations of nucleocytoplasmic transport by providing the native constraints on the diameter and a more precise mapping of the positions where the FG tails emanate from the scaffold (Fig. S14).

How an intricate structure consisting of ∼1000 components can be faithfully assembled in the crowded cellular environment is a very intriguing question. Our connectivity map captures the 3D trajectory of linker Nups through the assembled scaffold. Taken together with previous analysis of NPC assembly (*9*–*13, 19*), it suggests that the linker Nups facilitate dedicated spatial organization functions. The connections of Nup93 within individual IR complexes and to the Nup214-complex suggest a role in ensuring isostoichiometric assembly. This finding is consistent with the recent analysis of early NPC biogenesis, suggesting that Nup93 associates isostoichiometrically with the Nup62-complex already during translation in the cytosol (*58*). Thus, the stoichiometric assembly of the Nup62 subcomplex together with Nup205/188-Nup93 heterodimer is likely pre-assembled away from sites of NPC biogenesis, explaining the importance of the linker for intra-subcomplex interactions. How the spokes form a C2 symmetric interface at the NE plane remains to be addressed.

In the IR membrane coat, multiple interactions converge into an intriguing transmembrane interaction hub. We propose that its core is formed by Aladin-NDC1 heterodimer at the interface between the outer and inner rings. This transmembrane interaction hub is likely a spatial organizer for two proximate copies of Nup155 within the same spoke that point towards the outer rings and IR, respectively. Aladin-NDC1 likely further associates with Gp210, which arches between spokes in the NE lumen. The hub also binds Nup35, which connects to Nup155 copies of neighboring spokes, thus facilitating the horizontal, cylindrical oligomerization. Since Nup35 associates with Nup155 early during NPC assembly (*39*), its dimerizing domain appears critical to scaffold its flexible linkers towards neighboring spokes within IR membrane coat.

The often-emphasized notion that ‘NPCs fuse the INM and ONM’ or that they ‘stabilize the fusion of the INM and ONM’ is not necessarily supported by our analysis. Our simulations suggest that the membrane fusion topology *per se* is stable under certain conditions, relaxing toward a catenoid shape with zero membrane bending energy. Indeed, some species maintain the fusion topology in the absence of NPCs, e.g. during semi-closed mitosis in *Drosophila melanogaster* (*59*). Our analysis instead suggests that NPCs stabilize a pore that is wider than in the relaxed, tensionless double membrane hole. This notion agrees with the ultrastructural analysis of post-mitotic NPC assembly, which has revealed that NE holes are formed at small diameters and dilate once NPC subcomplexes are recruited (*60*). These data argue that the membrane shape defines the outline of the NPC scaffold and not *vice versa*.

A novel aspect of our work is that we use AI-based structure prediction programs AlphaFold and RoseTTAfold to model all atomic structures used for fitting to the EM maps and modeled the entire NPC scaffold without directly using any X-ray structures or homology models. Predicted atomic structures traditionally exhibited various inaccuracies, limiting their usage for detailed near-atomic model building in low-resolution EM maps. However, AlphaFold and RoseTTAfold have recently demonstrated unprecedented accuracy in predicting structures of monomeric proteins (*27, 28, 61*–*66*) and complexes (*35, 62, 67, 68*). The programs were also shown to accurately assess their confidence at the level of individual residues and inter-domain contacts (*27, 28, 62*). Indeed, we could successfully validate our models by comparing them to unpublished crystal structures, our cryo-EM maps, and biochemical data. The resulting model of the NPC scaffold is almost complete and exhibits near-atomic level precision at several interfaces. The model also contains several peripheral Nups, e.g. parts of the Nup214 and Nup358 complexes. Although the entire EM density for those peripheral Nups is unlikely to be resolved in the near future due to their flexibility, the complete model of the human NPC could be in reach by integrating data from complementary techniques that can address flexible proteins, such as super-resolution microscopy, FRET, and site-specific labeling (*18*).

Thanks to *in situ* and *in cellulo* cryo-ET and powerful AI-based prediction (*27, 28*), intricate structures such as the NPC can be now modeled. Not all subunit or domain combinations that we attempted to model with AI-based structure prediction led to structural models that were consistent with complementary data, emphasizing that experimental structure determination will still be required in the future for cases in which a priori knowledge remains sparse. However, even if AI-based modeling does not yield high confidence results, the models can still serve as tools for hypothesis generation and subsequent experimental validation.

## Acknowledgments

We acknowledge support from the Electron Microscopy Core Facility (EMCF) and IT services of EMBL Heidelberg.

## Funding

MB acknowledges funding by EMBL, the Max Planck Society, and the European Research Council (ComplexAssembly 724349). JK acknowledges funding from the Federal Ministry of Education and Research of Germany (FKZ 031L0100). M.S. and G.H.’s work on computer simulations was supported by the Max Planck Society. M.S. was supported by the EMBL Interdisciplinary Postdoc Programme under Marie Curie COFUND actions. M.S. and G.H. were supported by the Landes-Offensive zur Entwicklung Wissenschaftlich-ökonomischer Exzellenz (LOEWE) DynaMem program of the State of Hesse.

## Author contributions

SM prepared the samples, collected and analyzed the HeLa envelope data. SM, WH, and EM collected the HeLa envelope and HeLa *in cellulo* data. SM and BT performed the cryo-ET analysis. CEZ prepared, collected and analysed the HEK control and GP210Δ data. MTM performed the BioID experiments. FS and KB prepared the HEK GP210Δ cell line. AOK performed modeling and prepared figures, MS performed the MD simulations, GH performed the membrane elastic theory analysis, JK and AOK prepared software, SM, GH, MB, and JK conceptualized the study, supervised the project, and wrote the manuscript.

CRediT roles:

Conceptualization: SM, AOK, GH, MB, JK

Methodology: SM, AOK, BT, MS, JK

Investigation: SM, AOK, MS, BT, CEZ, KB, FS, EM, MTM, WH, GH, MB, JK

Visualization: SM, AOK, MS, CEZ, MTM, JK

Funding acquisition: MB

Project administration: SM, GH, MB, JK

Supervision: SM, GH, MB, JK

Writing – original draft: SM, MS, GH, MB, JK

Writing – review & editing: SM, AOK, MS, GH, MB, JK

## Competing interests

Authors declare that they have no competing interests.

## Data and materials availability

Prior publication, EM maps associated with the manuscript will be deposited in the Electron Microscopy Data Bank (EMDB) and the models will be deposited into the Protein Data Bank (PDB).

## Notes

### Competing Interest Statement

The authors have declared no competing interest.

### Summary of Updates

Fixed author name and affiliation, manuscript file did not change.

