## Supplementary Materials for "Artificial intelligence reveals nuclear pore complexity"

#### Supplementary Text

##### Pore shape from Helfrich membrane elastic theory

According to Helfrich membrane elastic theory, a double-membrane pore will relax to a catenoid shape, $r(z) = R\cosh(z/R)$, where $r$ and $z$ are radial and axial cylindrical coordinates, and $R$ is the pore radius. For a catenoid, the mean curvature is zero and the bending energy vanishes. In MD simulations of a double-membrane pore under periodic boundary conditions, we found the membrane shape to approach the catenoid (Fig. 4A). Further contraction in the simulations is limited by the distortions in a membrane of finite thickness and by membrane-membrane repulsion.

In the tomographic structures of the NPC, the double-membrane pore is substantially wider than expected for a catenoid. In the constricted state, the pore diameter is about $D = 2R \approx 90$ nm, and the distance between ONM and INM about $H \approx 30$ nm. We estimated the energy to deform the membrane to a diameter-to-width ratio of $x = D/H \approx 3$ using membrane elastic theory. We approximate the shape of the pore neck by a nodoid, which in cylindrical coordinates is given by

$$r(t) = a\left(\sqrt{\beta + cos^2 t} - cos\, t\right)$$

$$z(t) = a\left(sin\, t - \int_0^t \frac{cos^2 u}{\sqrt{\beta + cos^2 u}}\right)$$

for $-\pi/2 \leq t \leq \pi/2$, $a > 0$, and $\beta > 0$ with a pore radius of $R = r(0) = a\left(\sqrt{\beta+1} - 1\right)$. As a surface of constant mean curvature $H = 1/(2a)$, the nodoid is a minimum energy surface in membrane elastic theory for a non-zero spontaneous curvature or, equivalently, for a given area difference, as would be expected for an asymmetric membrane with protein anchors placed in the outer leaflet. The bending energy for the nodoid half-toroidal pore is

$$E_b = \frac{2\kappa A}{(2a)^2} = 2\pi\kappa\left[\sqrt{\beta}\left(2E(-1/\beta) - K(-1/\beta)\right) - 2\right]$$

with κ the membrane bending rigidity, and $K(u) = \int_0^{\pi/2} dt/\sqrt{1 - u\, sin^2 t}$ and $E(u) = \int_0^{\pi/2} dt\sqrt{1 - u\, sin^2 t}$ the complete elliptic integrals of the first and second kind. The ratio of pore diameter to inter-membrane spacing is

$$x = \frac{D}{H} = \frac{r(0)}{z(\pi/2)} = \frac{1 + \beta - \sqrt{1+\beta}}{\sqrt{1+\beta} - (1+\beta)E\left[\frac{1}{1+\beta}\right] + \beta K\left[\frac{1}{1+\beta}\right]}$$

For large $x = D/H$ ratios, the bending energy grows linearly as $E_b \sim \kappa(\pi^2 x - 10.45)$. For the experimentally observed ratios of $x \approx 3$, the bending energy of the nodoid-shaped pore is about $E_b \approx 20.5\kappa$. Even for the comparably low bending rigidity of $\kappa \approx 25$ kJ/mol reported for the nuclear envelope (*69*), this amounts to a bending energy of about 500 kJ/mol or about 200 $k_B T$. For reference, we also performed a variational minimization of the membrane bending energy with respect to the function $r(z)$ defining an axially symmetric pore shape in cylindrical coordinates. In addition to constraining the ratio $x$, we imposed infinite and zero slope at reduced arc lengths $\tau_0 = 0$ and $\tau_1 = 1$, respectively. For the minimization, we used a Fourier representation with 20 modes (*70*). The minimized bending energy is about $E_b \approx 18.2\kappa$, only about 10% lower than that for the nodoid. The energetic cost of widening the pore diameter to about three times the spacing between ONM and INM is thus substantial and requires compensation by protein-protein and protein-membrane interactions of the NPC that stabilize the pore shape.

### Materials and Methods

#### Mammalian cell cultivation

Modified human embryonic kidney cells 293 (HEK Flp-In$^{TM}$ T-Rex$^{TM}$) 293 Cell Line, Life Technologies) designed for rapid generation of stably transfected cell lines with a tetracycline-inducible expression system were used as parental cells. The GP210 CRISPR-knockout line (HEK GP210Δ) has been previously described (*71*). In general, all cells were maintained in Dulbecco's modified Eagle medium (DMEM) supplemented with 5g/L glucose and 10% heat inactivated fetal bovine serum (FBS, Sigma-Aldrich). HeLa Kyoto cell line was maintained in Dulbecco's modified Eagle medium (DMEM) medium containing 1 g/L glucose supplemented with 2 mM L-glutamine. Cells grown close to confluency (~90%) were trypsinized with 0.25% trypsin containing EDTA (Life Technologies) and passaged for further growth.

#### Grid preparation

Grids with HeLa nuclear envelopes were prepared exactly as described in (*21*). For the *in cellulo* work, Au200 R2/1 $SiO_2$ grids (Quantifoil Micro Tools GmbH) were glow discharged on both sides and sterilized under UV light. In a 6-well cell culture dish either 250,000 cells/well (for HeLa) or 400'000 cells/well (HEK293) were pipetted onto the grids pre-wetted with DMEM medium. Cells were left to settle and attach to the grids for 4h at 37°C in 5% $CO_2$. Subsequently the HeLa or HEK293 grids were plunge frozen with a Leica EM GP plunger with set chamber environment to 99% humidity and 37°C. Grids were blotted from the backside for 2 sec and plunged in liquid ethane-propane mix (37% and 63%) at ~ -195°C. HEK GP210Δ grids were washed once with PBS containing 8% dextran (35.45kDa) and blotted for 3 sec prior to plunge freezing in -186°C liquid ethane.

##### Cryo-FIB milling and Data acquisition

Plunge-frozen sample grids were FIB-milled on an Aquilos FIB-SEM (ThermoFisher Scientific) as described before (*22*, *23*). In brief, samples were coated with inorganic platinum (Pt-sputtering). Subsequently, a protective layer of organometallic platinum was deposited for ~ 20 sec using the gas injection system. Cells were then stepwise milled at a 20° angle to a final thickness of around 200-250 nm using decreasing ion-beam currents of 1 nA to 50 pA. A final round of Pt-sputtering was applied before unloading the sample.

##### Cryo-electron tomography and subtomogram averaging of the human NPC from nuclear envelopes

Tilt series were collected with SerialEM as described by Kosinski et al. (*19*). The angular coverage of the tilts spanned from -60 to +60 degrees. 10 8K x 8K frames, per tilt, were collected in the super-resolution mode on a K2 direct electron detector (Gatan Inc.) equipped with an BioQuantum Imaging Filter (GIF). An average total dose of 120 e-/Å2 per tomogram was used. 516 new tilt-series were collected and combined with 101 tilt-series reported by Kosinski et al. (*19*) leading to a total of 617 tilt-series. Incomplete tilt-series (missing more than 7 tilts in either direction or terminated due to autofocusing error because of the edge of the grid bar) were discarded. CTF was determined using CTFFind4 (*72*). Tilt-series with large discrepancies in the two defocus values estimated by CTFFind4 were also removed. This resulted in a total of 554 tilt-series. Tilt-series were manually aligned by tracking gold fiducials in IMOD (*73*). Tilt-series were filtered according to accumulated exposure based on parameters described by Kosinski et al. (*19*). Tilt-series were reconstructed with 3D CTF correction using NovaCTF (*74*).

7711 subtomograms containing individual NPCs were extracted, which correspond to 61688 asymmetric units. The pixel size at the specimen level was 3.37 Å. Tomograms were binned by Fourier cropping 2x (bin2), 4x (bin4) and 8x (bin8) and subtomograms were extracted at each level of binning corresponding to a pixel size of 6.74, 13.48 and 26.96 respectively. Subtomogram averaging was performed on a whole pore level with bin8 dataset. Subsequently, asymmetric units were extracted from the aligned pores and averaged as described in Kosinski et al. (*19*). Subunits with the center outside of tomogram boundaries were excluded from further processing. The cytoplasmic (CR), inner (IR) and nuclear (NR) ring were processed separately, i.e. the positions of subunit centers were moved to be in the center of each ring, resulting in three different set of subtomograms. Rings with the center outside of tomogram boundaries were excluded from further processing. The subtomograms were iteratively aligned first on bin4, then on bin2 level and the final alignment was refined on bin1 level. The complete subtomogram averaging and alignment was performed using novaSTA (*75*), the masks necessary for the alignment were created in Dynamo (*76*) and Relion (*77*).

After the bin4 alignment, the quality of each subtomogram was assessed using the expected geometrical shape of a complete ring. More precisely, the subunits corresponding to one NPC ring should still be part of a ring after the alignment. For each ring and each subunit, the angular distance of its normal vector to all other normal vectors of subunits within the same ring was computed. Subsequently, the distances were averaged (for each NPC separately) and for each subunit the deviation from the normal vector from the average was computed. The same was done for so-called in-plane vectors, i.e. vectors describing the direction from the NPC center to the center of a subunit. The expected/ideal angular distance for the normal vectors is zero while for the in-plane vectors it is 45°. This analysis was performed only on the rings with at least three subunits that were retained from the initial sub-tomogram averaging runs. The rings with less than three subunits were removed from further processing. The computed deviations were used to identify poorly aligned ring subunits in order to remove them. For CR and SR all subunits with the normal vector deviation greater than 30° were removed. The deviation of in-plane angles from expected 45° greater than 15° or CR and 10° for SR was used as threshold for additional removal of ring subunits. The threshold values were determined empirically. For CR and SR, the number of subunits left for processing after geometrical cleaning were 31774 and 35281 respectively. The final subunit cleaning to remove poor quality subunits was performed on the final average using the constrained cross-correlation (CCC) value which was computed between each subtomogram and the reference during the last iteration of alignment. Subtomograms with the worst CCC values were subsequently removed in batches of 1000 as long as the resolution improved. The number of subunits/subtomograms contributing towards the final structure of the CR and SR ring were 21604 and 30000, respectively.

In contrast to CR and IR, adding additional tilt-series followed by geometric cleaning procedure on NR did not yield any significant improvement in comparison to the dataset reported in Kosinski et.al (*19*). Thus, the original map of the NR was used for the presented analysis. The map was created using steps described in Kosinski et al. (*19*) and the total number of particles contributing to the final average were 11112.

##### Cryo-electron tomography and subtomogram averaging of human NPC *in cellulo*

Data acquisition was performed on a Titan Krios G2 (for HeLa) or G4 (for HEK) (ThermoFisher), operating at 300kV and equipped with Gatan K2 Summit direct electron detector and energy filter as described before (*23*). In brief, tilt series were acquired in dosefractionation mode at 4k x 4k resolution with a nominal pixel size of 3.37 (HeLa) or 3.45 Å (HEK) using a automated dose-symmetric acquisition scheme (*78*) starting at a given pre-tilt corresponding to the tilt of the FIB-milled lamellae (typically +/- 13°). Tilt series were acquired with a tilt increment of 3° and a tilt range interval of -50°/+50°, and a total dose per tomogram of 120-150 e/Å$^2$.

Tilt series preprocessing and tomogram reconstruction was performed as described previously (*22*, *23*). Subtomogram averaging was performed as described before (*22*, *23*) . In brief, for the HeLa control dataset 53 NPCs were extracted from 13 tomograms. For the HEK dataset, 30 control and 43 gp210Δ NPCs were extracted form 8 control and 14 gp210Δ tomograms respectively. Whole pores were aligned using bin8 and bin4 subtomograms with imposed 8-fold symmetry. Upon convergence 280, 150 and 222 subunits were extracted from the control HeLa, control HEK and HEKGp210Δ data set respectively and the cytoplasmic, inner and nuclear ring (CR, IR and NR) subunits were further refined independently using bin4 subtomograms. The individual ring subunits were refined without splitting the data into independent half sets to a final resolution of <54 Å (NR) and <48 Å (CR and IR) as estimated by Fourier Shell Correlation (FSC) using the 0.5 criterion for the HEK datasets. For the HeLa dataset gold-standard criteria was used to calculate the FSC, which resulted in final resolution of 45 Å (CR and IR) and 53 Å for the NR.

##### Proximity labeling using BioID

BioID analysis of Aladin was done as previously described (*79*). In brief, Aladin was BirA tagged and over-expressed in Hek293 Flp-In Trex cells. Quantitative mass spectrometry was done in four biological replicates and in comparison to control cells expressing BirA tagged NLS-NES-Dendra that resides within the central channel.

##### Structural modeling of Nups and NPC subcomplexes

The structures of all individual Nups and selected subcomplexes were modeled using AlphaFold (*27*) or downloaded from AlphaFold Database (*61*). The models of monomeric proteins (Nup155, Nup133, Nup107, Nup93, Nup205, Nup188, Nup160, Nup358 and Elys), were download from from AlphaFold Database (*61*). To model subcomplexes or their parts around the interfaces (Nup62-Nup54-Nup58, Nup205-Nup93, Nup188-Nup93, Nup155-Nup53, Nup93-Nup53, NDC1-Aladin, Nup35 homo-dimer, Nup85-Seh1-Nup43, Nup160-Nup96-Sec13, Nup160-Nup37, Nup133-Nup107, Nup96-Nup107, and Nup160-Nup96-Nup85, Nup214-Nup62-Nup88, Nup88-Nup98), we used the AlphaFold version modified for modeling complexes, available through ColabFold (*36*) with all parameters set to default except max_recycles parameter set to between 12 and 48 depending on the subcomplex. For Gp210, we first built the initial full-length using RoseTTAfold (*28*), as AlphaFold did not provide a full-length model fitting well into the EM density map. After fitting the model into the EM maps as a rigid body (see below), we used AlphaFold to model successive monomeric and homodimeric fragments of Gp210, superposed them onto the fitted RoseTTAfold model, and refined the fits. The quality of the AlphaFold models was first assessed by the scores provided by the authors—the predicted local-distance difference test (pLDDT), which predicts the local accuracy, and Predicted Aligned Error, which assesses the packing between domains and protein chains. In addition, we validated the models by comparing to structures not used for modeling, yet unpublished structures provided in the accompanying paper, fits to the cryo-ET maps, and previously published biochemical data (Fig. S5-Fig**. S9**, Fig. S11).

#### Systematic fitting of atomic structures to cryo-ET maps

We used the previously published procedure for systematic fitting (*8*, *19*, *21*, *22*, *26*, *37*, *80*) to both locate the atomic structures in the cryo-ET maps and validate the AlphaFold models. Prior to fitting, all the high-resolution structures were filtered to 10-15 Å. The resulting simulated model maps were subsequently fitted into individual ring segments of cryo-ET maps by global fitting as implemented in UCSF Chimera (*81*) using scripts within Assembline (*37*). The maps used for fitting did not include nuclear envelope density in order to eliminate the possibility of fits significantly overlapping with the membrane. All fitting runs were performed using 100,000 random initial placements with the requirement of at least 30-60% (depending on the size of the structure) of the simulated model map to be covered by the cryo-ET density envelope defined at a low threshold. For each fitted model, this procedure resulted in around 1000-20,000 fits with non-redundant conformations upon clustering. The cross-correlation about the mean (cam score, equivalent to Pearson correlation) score from UCSF Chimera (Pettersen et al., 2004) was used as a fitting metric for each atomic structure, similarly to our previously published works. The statistical significance of every fitted model was evaluated as a p-value derived from the cam scores. The calculation of p-values was performed by first transforming the cross-correlation scores to z-scores (Fisher's z-transform) and centering, from which subsequently two-sided p-values were computed using standard deviation derived from an empirical null distribution (based on all obtained non-redundant fits and fitted using fdrtool (*82*) R-package). Finally, the p-values were corrected for multiple testing with Benjamini-Hochberg (*83*).

#### Modeling of the human NPC scaffold

To assemble the models of the entire NPC scaffold based on the constricted and dilated cryo-ET maps we used our integrative modeling software Assembline (*37*), which is based on Integrative Modeling Platform (IMP) (*84*) version 2.15 and Python Modeling Interface (PMI) (*85*). First, we built the model of the constricted NPC owing to its higher resolution. The AlphaFold models of Nup domains and sub-complexes already present in our previous human NPC models (*19*, *21*), were placed in the map by superposing them onto the published models. The remaining domains and sub-complexes added in this work (Nup358, Nup53, Nup93 in the outer rings, NDC1 and Aladin, Nup214 complex), were placed using systematic fitting (as above) and global optimization procedure of Assembline. In addition to using models of sub-complexes as rigid bodies for fitting, several inter-subunit interfaces were restrained by elastic distance network derived from ColabFold models overlapping with and bridging already fitted models. During the refinement, the structures were used as rigid-bodies and simultaneously represented at two resolutions: in Cα-only representation and a coarse-grained representation, in which each 10-residue stretch was converted to a bead. The 10-residue bead representation was used for all restraints to increase computational efficiency except for the domain connectivity restraints, for which the Cα-only representation was used. The flexible protein linkers between the domains were added as chains of one-residue beads. The entire structure was optimized using the refinement step of Assembline to optimize the fit to the map, minimize steric clashes, and ensure connectivity of the protein linkers. The scoring function for the refinement comprised the EM fit restraint, clash score (SoftSpherePairScore of IMP), connectivity distance between domains neighboring in sequence, a term preventing overlap of the protein mass with the nuclear envelope, a restraint promoting the membrane-binding loops of Nup133, Nup160, Nup155 to interact with the envelope, implemented using MapDistanceTransform of IMP (predicted by similarity to known or predicted ALPS motifs *X. laevis* and *S. cerevisiae* homologs (*6*, *21*, *86*), and elastic network restraints derived from the sub-complexes modeled with AlphaFold/Colabfold. The final atomic structures were generated based on the refinement models by back-mapping the coarse-grained representation to the original AlphaFold atomic models. The conformation of the linkers was further optimized using Modeller (*87*) and Isolde (*88*).

The model of the dilated NPC was built by fitting the asymmetric units of the individual cytoplasmic, inner and nuclear rings of the constricted NPC model to the dilated cryo-ET maps and refining the fits with Assembline. The refinement procedure was performed as above.

To calculate the percentage of the molecular weight of the full NPC and the NPC scaffold covered by the new and the old models, we defined the full NPC as composed of the following 32 Nups, with the stoichiometry indicated in the parentheses: Nup160 (32), Nup96 (32), Nup85 (32), Seh1 (32), Sec13 (32), Nup107 (32), Nup133 (32), Nup358 (40), Nup43 (32), Elys (16), Nup37 (32), Nup188 (16), Nup205 (40), Nup155 (48), Nup93 (56), Nup53 (32), Nup62 (48), Nup54 (32), Nup58 (32), Nup88 (16), Nup214 (16), Nup98 (48), NDC1 (16), NUP210 (64), and Aladin (16), POM121 (32), TPR (32), NUP153 (32), NUP50 (16), CG1 (8), DDX19 (16), GLE1 (8). The scaffold NPC was defined as composed of 25 Nups: Nup160, Nup96, Nup85,

Seh1, Sec13, Nup107, Nup133, Nup358, Nup43, Elys, Nup37, Nup188, Nup205, Nup155, Nup93, Nup53, Nup62, Nup54, Nup58, Nup88, Nup214, Nup98, NDC1, NUP210, and Aladin. The stoichiometry for the scaffold was the same as for the full NPC with exception of Nup214 complex for which only one copy was counted, as the second copy is not clearly visible in the EM density. Note that for some nucleoporins like Nup98 or POM121, the exact stoichiometry is still uncertain. The coiled-coil domains of the peripheral Nups of the Nup214 complex and the $\alpha$-solenoid domain of Nup358 were included in the scaffold. The FG regions were excluded. These definitions resulted in the molecular weight of 119 MDa for the full NPC and 76 MDa for the scaffold. The scaffold diameters were described by two distances between the opposite spokes: the membrane-to-membrane distance and the distance between ferredoxin-like domains of Nup54 at the residue 220.

Figures were produced using UCSF ChimeraX (*89*).

##### Molecular dynamics (MD) simulations

All molecular dynamics simulations were performed using the GROMACS software package and the coarse-grained Martini force field v2.2 (*54*, *90*, *91*).

Toroidal membrane pores were constructed by building a double bilayer cylindrical shape with the BUMPy tool (*92*). A pre-equilibrated coarse-grained POPC lipid bilayer was used as input, which was generated using *insane.py* (*92*, *93*). The toroidal membrane profile was manually matched to the membrane density from cryo-ET. Two carbon nanotube porins (CNTPs) were inserted into the membrane to enable solvent transfer between the two separated volumes, as is required for membrane-mechanical equilibration. CNTPs were built according to previous work (*92*, *93*). The generated CNTPs had a length of 3.6 nm and a diameter of 14.7 nm. The two outermost rings consisted of SNda beads to enable stable membrane embedding (*94*). To increase stiffness of these large CNTPs the improper dihedral force constant was increased to 1000 kJ mol$^{-1}$ rad$^{-2}$. The CNTPs were embedded in the flat patch of the NPC membranes away from the NPC. Lipids within 8 Å of the CNTP or inside the circumference of the tube were removed. The CNTP parameters were used as previously reported (*94*, *95*).

All protein structures were coarse-grained using the *martinize.py* script. Each protein subunit was coarse grained individually. The model used for simulations did not include GP210 for simplicity. All chain termini were uncharged. Secondary structure restraints were assigned according to DSSP (*96*). The tertiary structure of the proteins was maintained by an elastic network with cutoff $R_c = 0.9$ nm and force constant $k = 1000$ kJ mol$^{-1}$ nm$^{-2}$ was applied to each protein. For all proteins, the ElNeDyn2.2 protein force field was used in conjunction with Martini 2.2 (*54*, *55*). Simulations were performed with the default protein-protein interactions $\alpha = 1.0$ (results shown in Supplementary Materials), as well as a scaled version with $\alpha = 0.7$

(*97*) (results shown in the main text) to test the effect of reported overestimated non-bonded interactions (*98*). The hNPC structure was centered within the membrane pore models containing the CNTPs and overlapping lipids were removed. All systems were solvated with coarse-grained water containing 10% anti-freeze WF particles and $Na^+$ ions to neutralize the system. All simulated systems used in this study are listed in **Table S1**.

Each system was steepest-descent energy minimized using a soft-core potential to remove steric clashes of lipids and proteins. The systems were then equilibrated in an NPT ensemble with semisotropic pressure coupling first for 2.5 ns with a 5 fs timestep and then for 100 ns with a 15 fs timestep with position restraints on the protein backbone beads with a force constant of 1000 kJ $mol^{-1}$ $nm^{-2}$, maintaining a temperature of 310 K and pressure of 1 bar using the Berendsen barostat and velocity rescaling thermostat (*99*, *100*). Characteristic coupling times of 12 and 1 ps were used, respectively. During production simulations the Parrinello-Rahman barostat was used (*101*).

To apply lateral tension, an anisotropic pressure tensor was used with an out-of-plane pressure of $P_\perp = p + 2\Delta P/3$ and an in-plane pressure of $P_\parallel = p - \Delta P/3$, with p = 1 bar. This results in a traceless lateral strain $S = \mathrm{diag}(-\Delta P/3, -\Delta P/3, 2\Delta P/3)$ where $\Delta P \equiv P_\perp - P_\parallel$. The resulting tension on the double-membrane system is $\sigma = (P_\perp - P_\parallel)L_z = \Delta P L_z$ with $L_z$ the box height. To allow for gradual equilibration under tension, ΔP was increased in steps of 1 bar until reaching the target value (see Table S1).

The Verlet neighbor search algorithm was used to update the neighbor list, with the length and update frequency being automatically determined. Lennard-Jones and Coulomb forces were cut off at 1.1 nm with the potential shifted to 0 using the Verlet-shift potential modifier. A 15 fs timestep was used in all production simulations. Production simulations were performed for 1 μs each.

Images and movies were generated using VMD and time series were analyzed using the MDAnalysis library (*102*, *103*).

To calculate the root-mean-square distance (RMSD) we superimposed the structures onto the structure from the initial model using the qcprot RMSD alignment algorithm implemented in MDAnalysis (*103*). The RMSD was calculated every 1.5 ns for the BB beads with respect to the rigid-body aligned initial structure. As units, each individual protein was considered as well as each of the eight spokes as a whole. Averages and standard deviations were calculated from the eight spokes or all equivalent protein copies, respectively.

The diameter of the NPC pore was analyzed by fitting a circle to the membrane midplane (C4A, and C4B beads) at the narrowest point of toroidal NPC membrane in the xy-plane.

We note that the time scale of the simulations that can be computed is too short to recapitulate the complete NPC dilation and constriction processes. In addition to mechanical tension in the widened double-membrane pore (Supplementary Notes), we expect the missing FG mesh in the MD model to contribute to the compaction of the NPC scaffold seen in the MD simulations. We also note that the elastic network on proteins of the Martini model restricts internal conformational changes, which might be required for larger scale NPC dilation.

#### Supplementary Tables

**Table S1 Overview of MD simulations.**

| **Protein** | **System size [nm$^3$]** | **No. of particles** | **Alpha scaling** (protein-protein) | **Pxx=Pyy, Pzz [bar]** (after equil) | **Simulation time** |
|---|---|---|---|---|---|
| Without hNPC | 140×140×80 | 13813071 | 1.0 | 1, 1 | 1 μs |
| With hNPC | 140×140×80 | 13395936 | 1.0 | 1, 1 | 1 μs |
| With hNPC | 140×140×80 | 13395936 | 0.7 | 1, 1 | 1 μs |
| With hNPC | 140×140×80 | 13395936 | 1.0 | 0.33,2.33 | 1 μs |
| With hNPC | 140×140×80 | 13395936 | 0.7 | 0.33,2.33 | 1 μs |
| With hNPC | 140×140×80 | 13395936 | 1.0 | 0, 3 | 1 μs |

#### Supplementary Figures

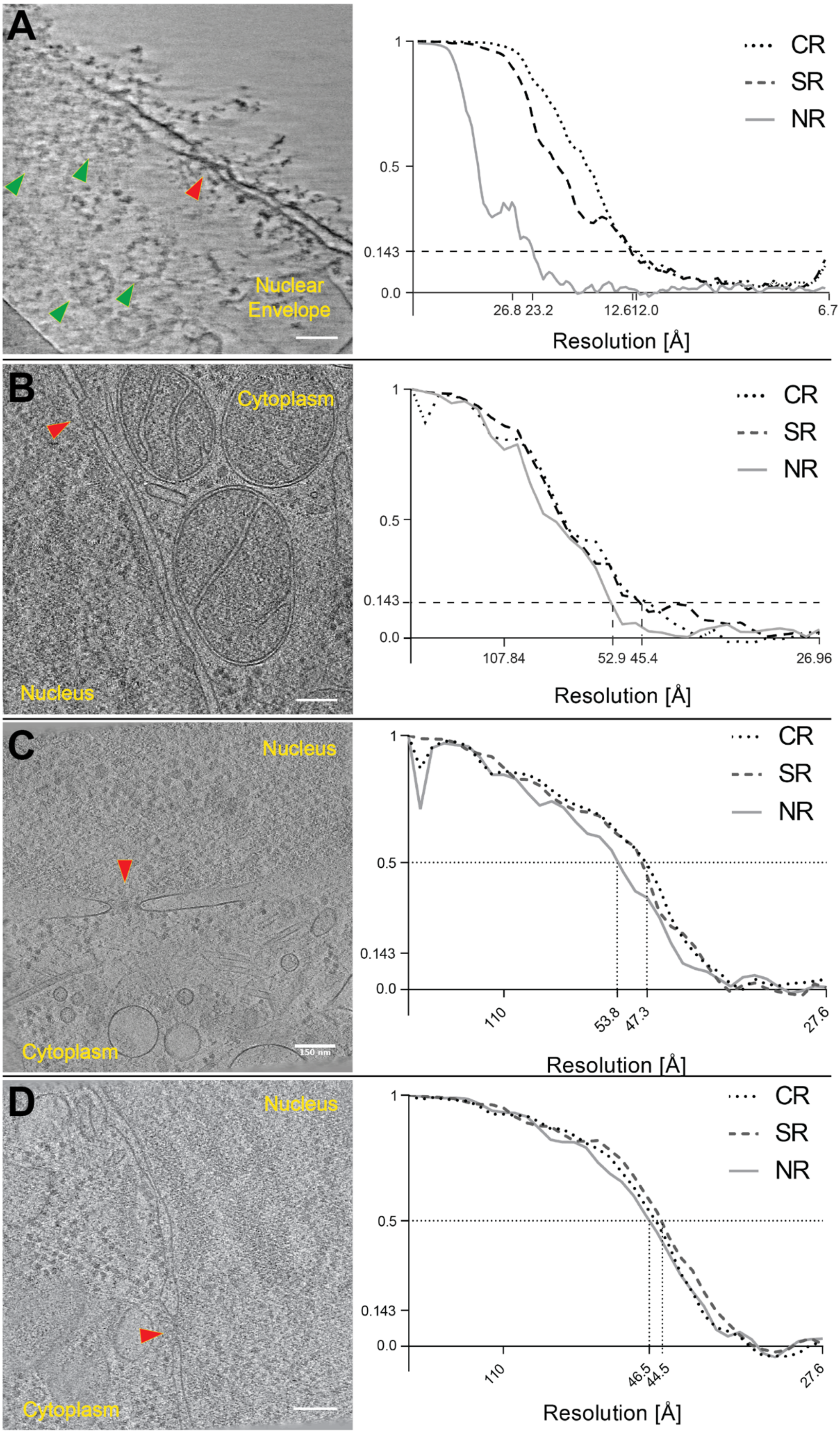

**Fig. S1 Slices through tomograms and corresponding Fourier Shell correlation plots.** Slices through representative tomograms of isolated nuclear envelopes **(A)**, cryo-FIB milled HeLa cells **(B)**, cryo-FIB milled HEK cells **(C)** or cryo-FIB milled HEKgp210Δ cells **(D)**. NPCs are highlighted with red arrowheads (side view) or green arrowheads (top view, seen only in case of nuclear envelopes). Fourier Shell Correlation plots of the corresponding NPC averages of the individual rings is shown on the right. Scale bar: 100 nm in (A), 150nm in (B-D).

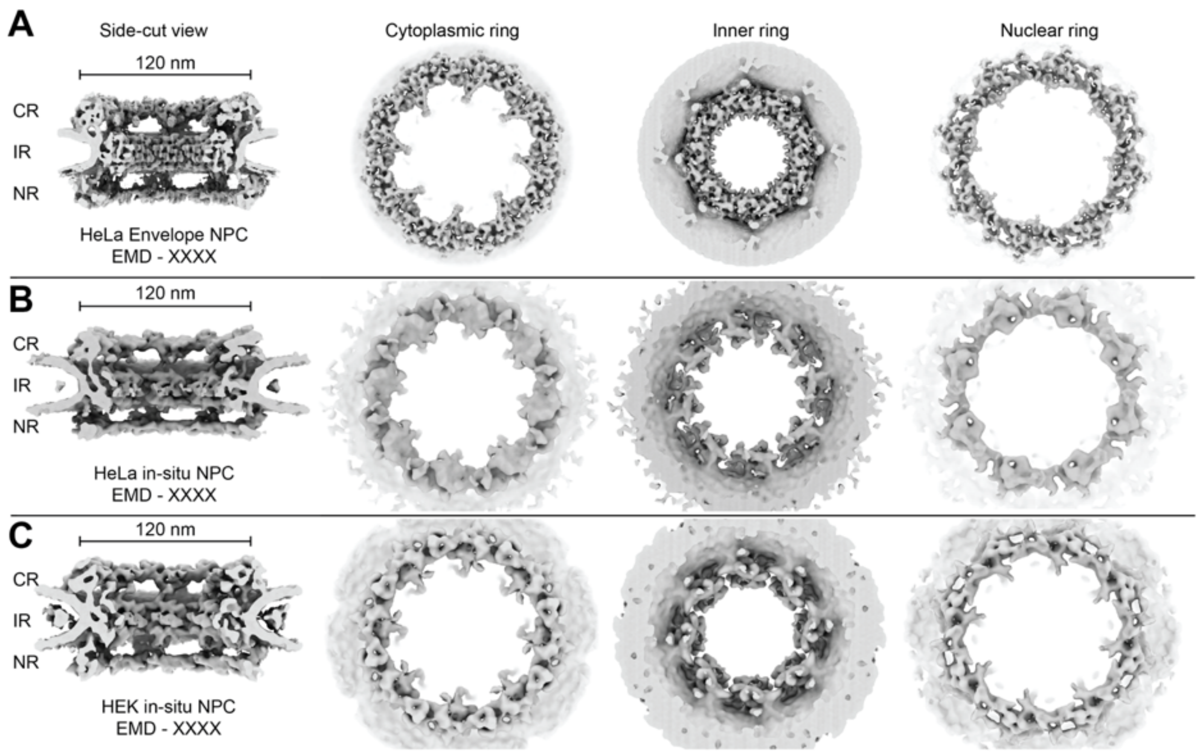


**Fig. S2 Structure of human NPCs from isolated NEs or *in cellulo.*** Isosurface views of the cut-side view, cytoplasmic, inner and nuclear rings of the structure of NPC obtained from isolated HeLa NE **(A)**, cryo-FIB milled control HeLa cells **(B)** and cryo-FIB milled control HEK cells **(C)** are shown (from L-R). The *In cellulo* NPCs have a larger central channel diameter in comparison to NPC from isolated HeLa envelope.

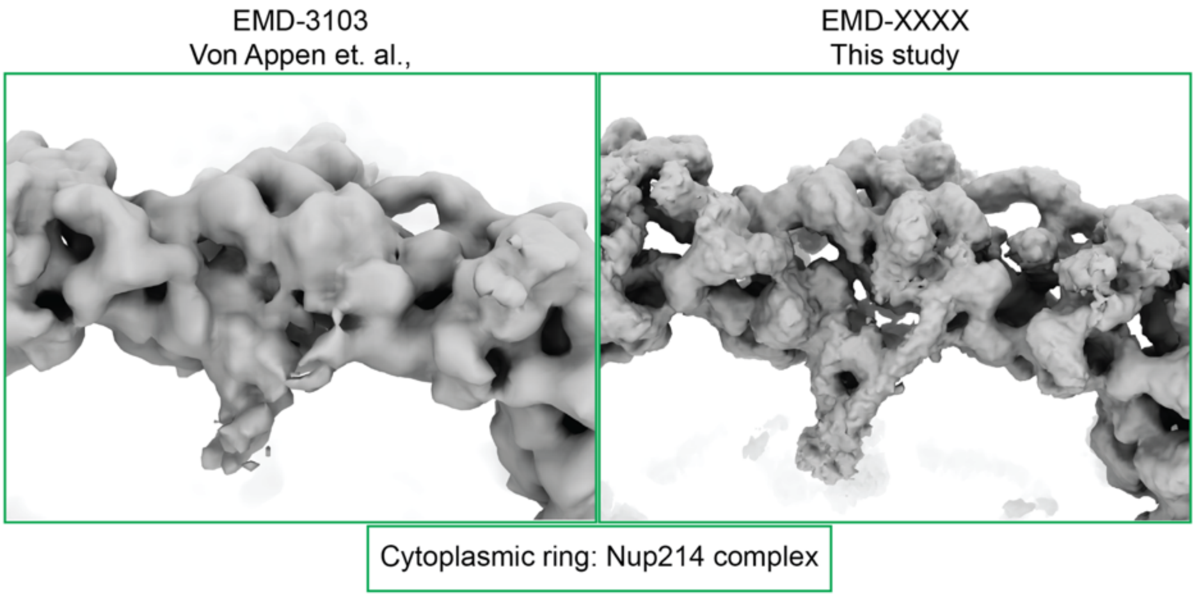


**Fig. S3 Comparison of human NPC structures.** Resolution improvement in our new structure of NPC from isolated HeLa envelopes (right) in comparison to existing human structure (left). Region of the cytoplasmic ring containing Nup214 subcomplex is highlighted.

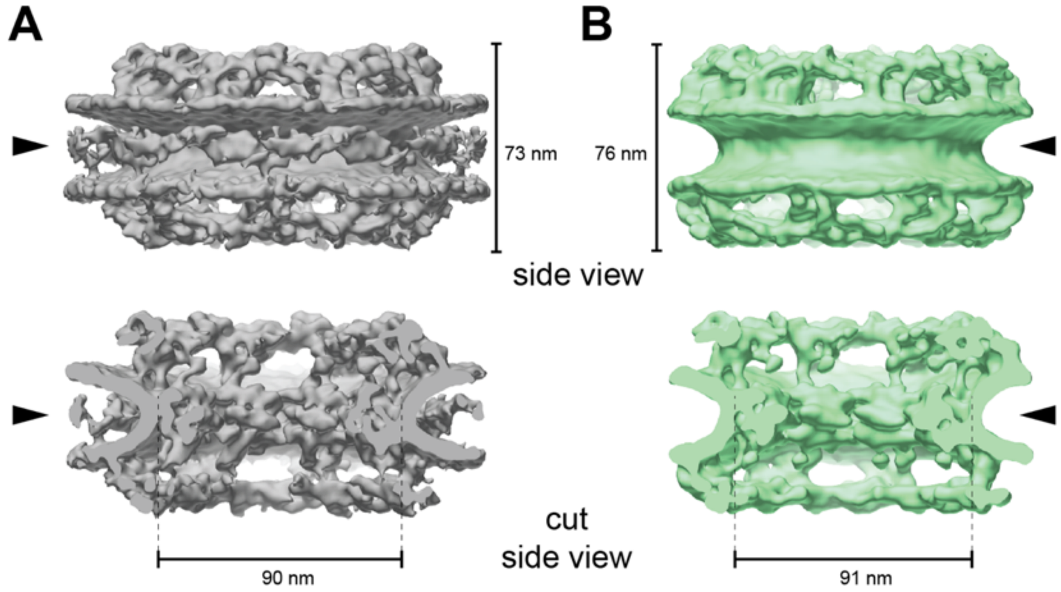


**Fig. S4** ***In cellulo*** **human NPC cryo-ET maps from HEK control and gp210Δ cells.** Isosurface representation non-modified (grey, left) and gp210Δ (green, right) human *in cellulo* NPC cryo-EM maps shown in side view **(A)** and cut-side view **(B)** along the central axis. Black arrowheads highlight the luminal density corresponding to Gp210 which is present in the control (left) and missing in the gp210Δ map.

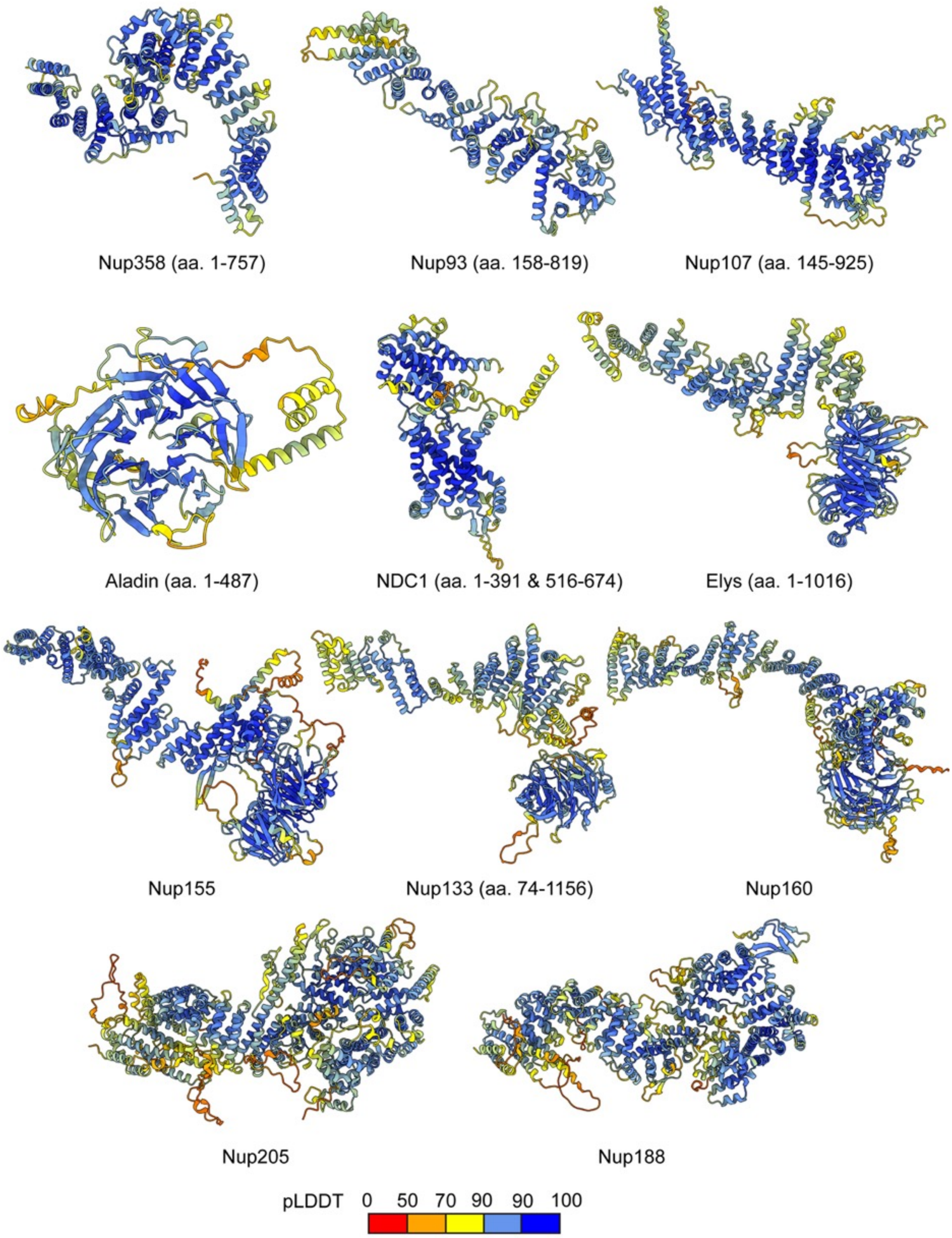
Nup358 (aa. 1-757)
Nup93 (aa. 158-819)
Nup107 (aa. 145-925)
Aladin (aa. 1-487)
NDC1 (aa. 1-391 & 516-674)
Elys (aa. 1-1016)
Nup155
Nup133 (aa. 74-1156)
Nup160
Nup205
Nup188
pLDDT 0 50 70 90 90 100

**Fig. S5 Estimated quality of monomeric structural models of Nups built using AlphaFold.** The models are colored by local model confidence estimated with predicted local distance difference test (pLDDT), as returned by AlphaFold. The pLDDT > 90 (dark blue) indicates high accuracy of backbone and side chain rotamers whereas pLDDT > 70 (yellow) indicates correct backbone prediction (*61*). Long low confidence regions are hidden from view for clarity. aa. – amino acid residues.

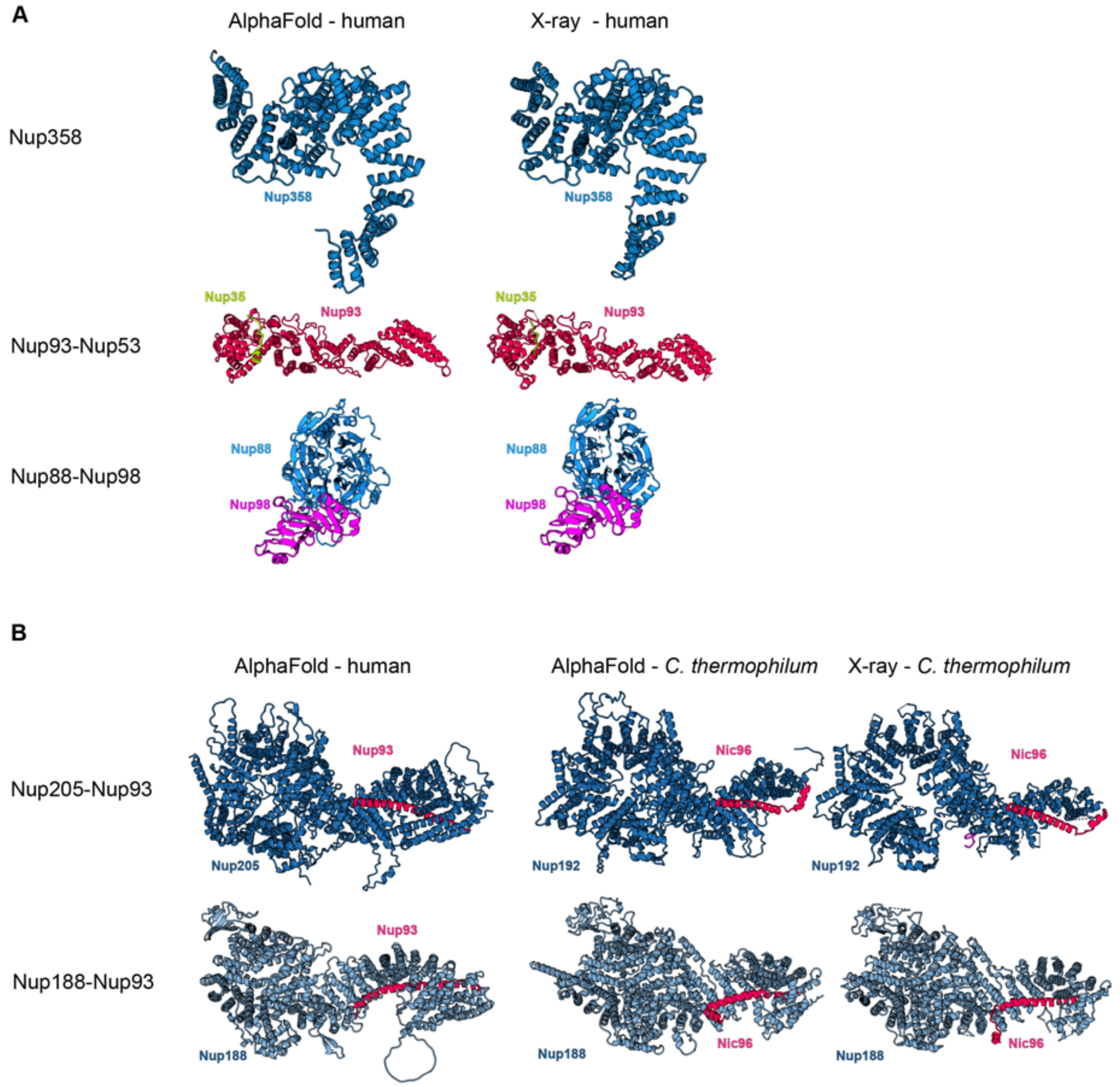


**Fig. S6 Validation of selected AlphaFold models with X-ray crystallography structures.** The models are compared to X-ray crystallography structures that have been released in the accompanying work (to be published and referenced here upon acceptance) and have not been used as templates for modeling. **(A)** Models are compared to structures of sub-complexes from human. **(B)** Models of Nup205-Nup93 and Nup188-Nup93 complexes are compared to X-ray structures from *C. thermophilum*, as X-ray structures for human are not available. The structures exhibit some differences due to inter-species variation but harbor the same interaction sites. For comparison, AlphaFold models of *C. thermophilum* are also shown, and recapitulate the X-ray structures. Proteins are colored as in Fig. 1.

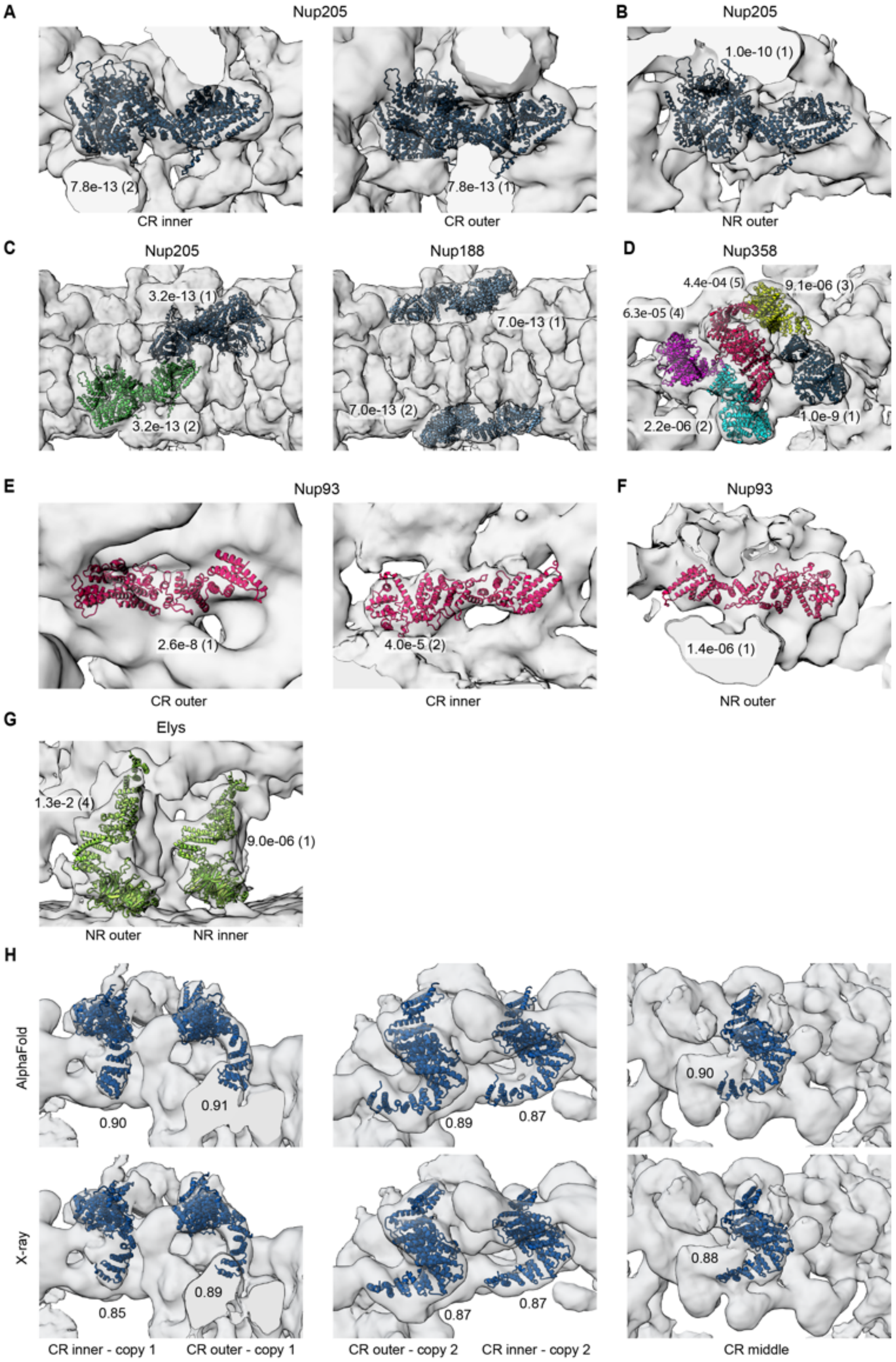


**Fig. S7 Systematic fitting of monomeric AlphaFold models of Nups previously not included in the human NPC model or till now not localized unambiguously in the human NPC.** Each panel shows the top fits fitted to the EM maps. P-values, calculated as described in the Methods, and ranks of the fits are indicated next to the structures. **(A)** Nup205 fitted to the map of the CR assigned two copies of Nup205 in the CR **(B)** Nup205 fitted to the map of the NR assigned one copy of Nup205 in the NR **(C)** Nup188 and Nup205 fitted to the map of the IR assigned Nup188 to the outer sub-complexes of the IR and Nup205 to the inner sub-complexes of the IR. **(D)** Nup358 was fitted to the EM map of the entire CR spoke assigning five fits, and thus five copies, of Nup358 per NPC spoke (*left*). **(E and F)** Nup93 fitted to the difference maps of the CR and NR obtained after subtracting the density of Y complexes identifies two locations of Nup93 in CR and one in the NR. **(G)** Nup93 fitted to the difference map of the NR confirms two copies of Elys in the NR. (H) AlphaFold model of Nup358 fits the EM density better than the Nup358 X-ray structures (only the better fitting of the two X-ray structure is shown). Numbers next to the fits indicate cross-correlation with the EM density as calculated with UCSF Chimera.

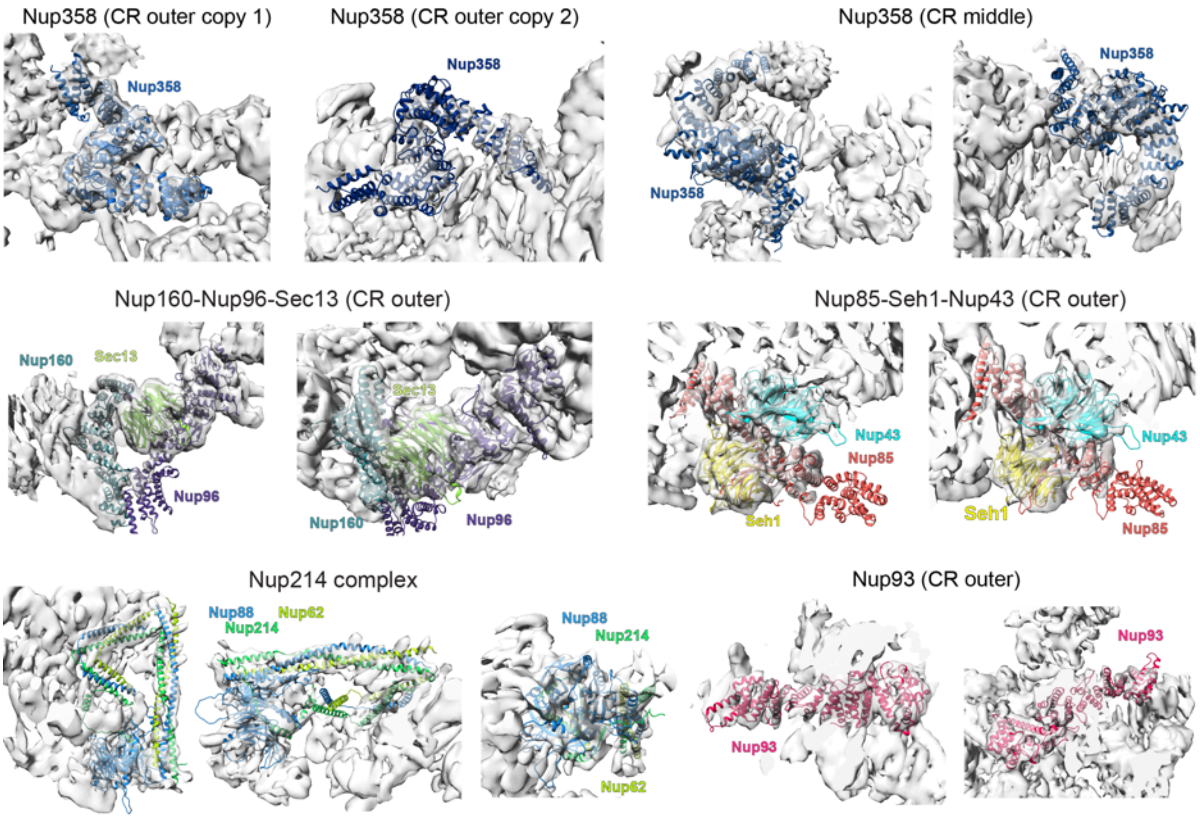


**Fig. S8 Consistency of secondary structure observed in the EM map of *Xenopus laevis* CR subunit (*29*) with AI-based models of human Nups at the respective fit positions into the human NPC.** The components of the CR subunit modeled with AlphaFold are fitted to the EM map of *Xenopus laevis* CR subunit (EM Data Bank ID EMD-0910) and EMD-0911 (for Nup214 complex). The quality of the AlphaFold models can be appreciated by the match to secondary structure densities in the EM maps. Proteins are colored as in Fig. 1.

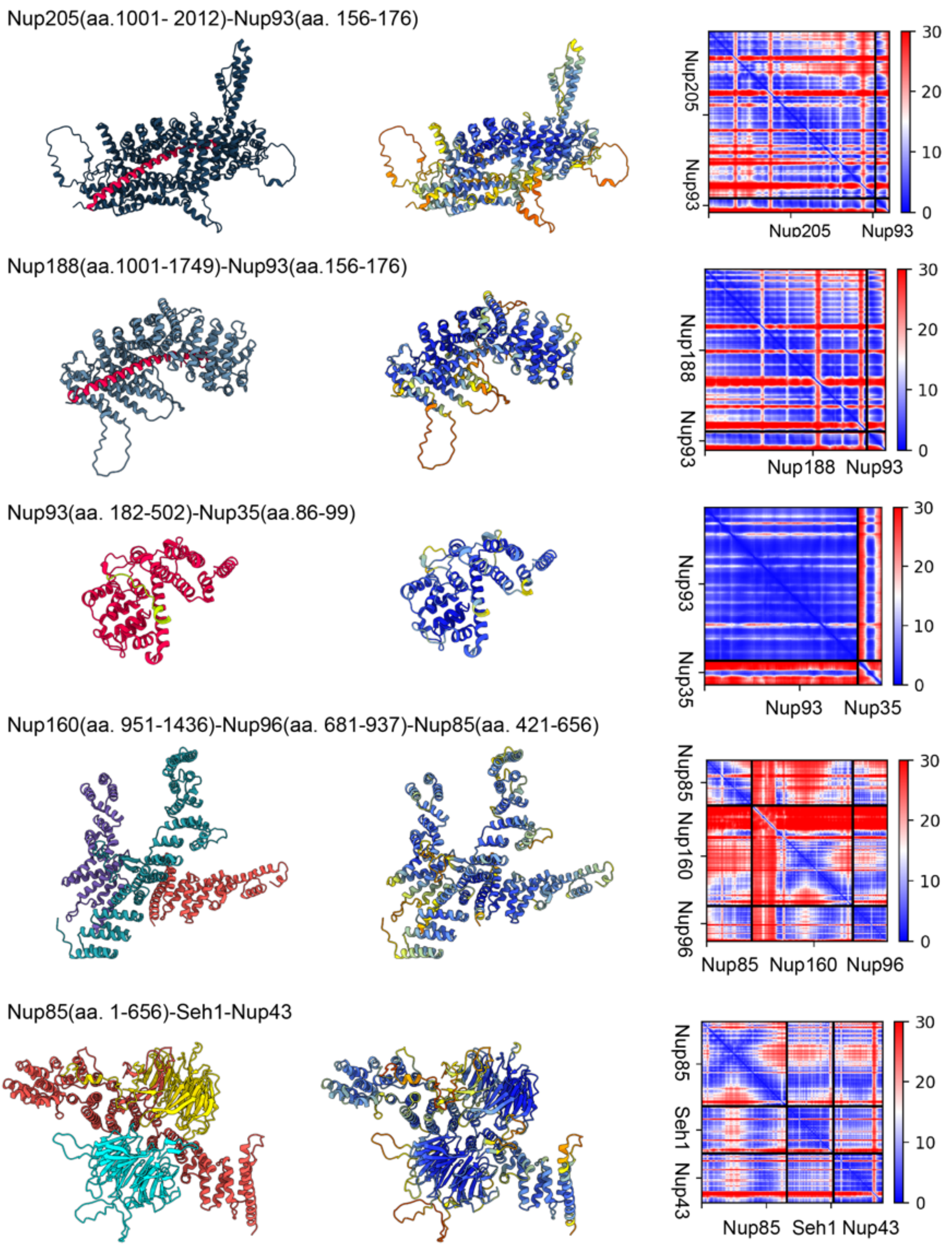
Nup205(aa.1001- 2012)-Nup93(aa. 156-176)
Nup205
Nup93
Nup205
Nup93
30
20
10
0
Nup188(aa.1001-1749)-Nup93(aa.156-176)
Nup188
Nup93
Nup188
Nup93
30
20
10
0
Nup93(aa. 182-502)-Nup35(aa.86-99)
Nup93
Nup35
Nup93
Nup35
30
20
10
0
Nup160(aa. 951-1436)-Nup96(aa. 681-937)-Nup85(aa. 421-656)
Nup85
Nup160
Nup96
Nup85
Nup160
Nup96
30
20
10
0
Nup85(aa. 1-656)-Seh1-Nup43
Nup85
Seh1
Nup43
Nup85
Seh1
Nup43
30
20
10
0

Nup214(aa. 1-428,700-1000)-Nup88-Nup62(aa. 332-502)

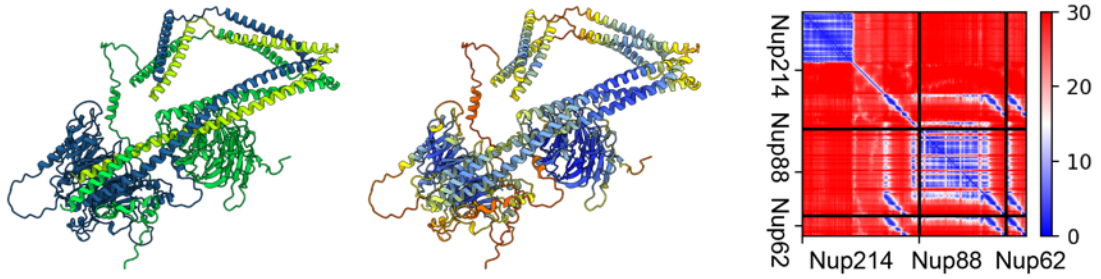


Nup62(aa. 332-502)-Nup58(aa. 246-418)-Nup54(aa. 111-493)-Nup93(aa. 1-94)

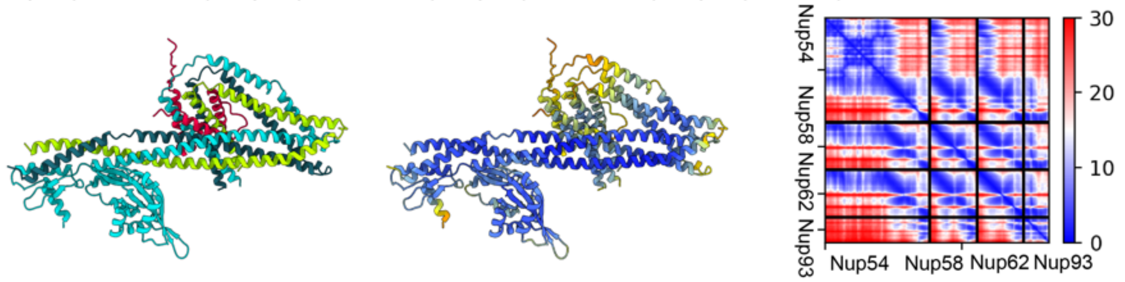


Nup98(aa. 730-880)-Nup88(aa. 1-579)

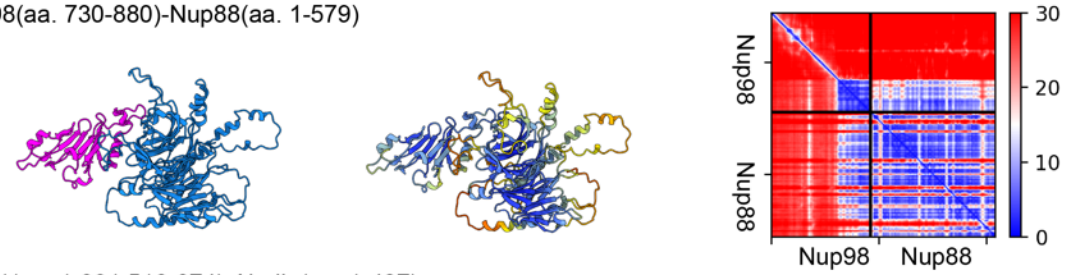


NDC1(aa. 1-391,516-674)-Aladin(aa. 1-487)

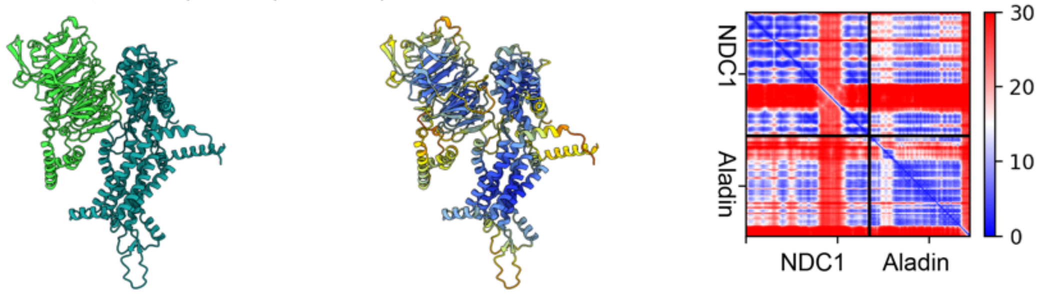


Nup155(aa. 40-577)-Nup35(aa. 285-326)

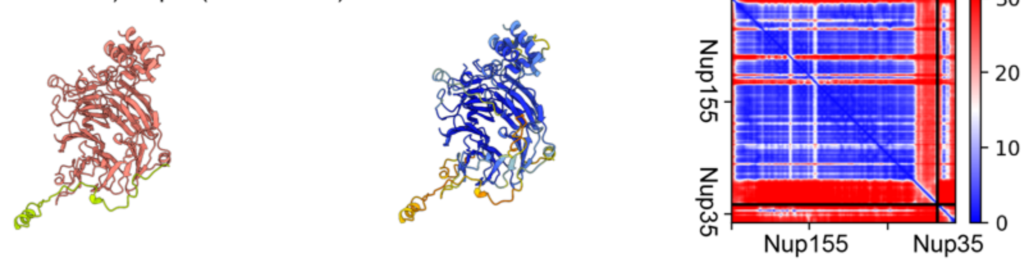

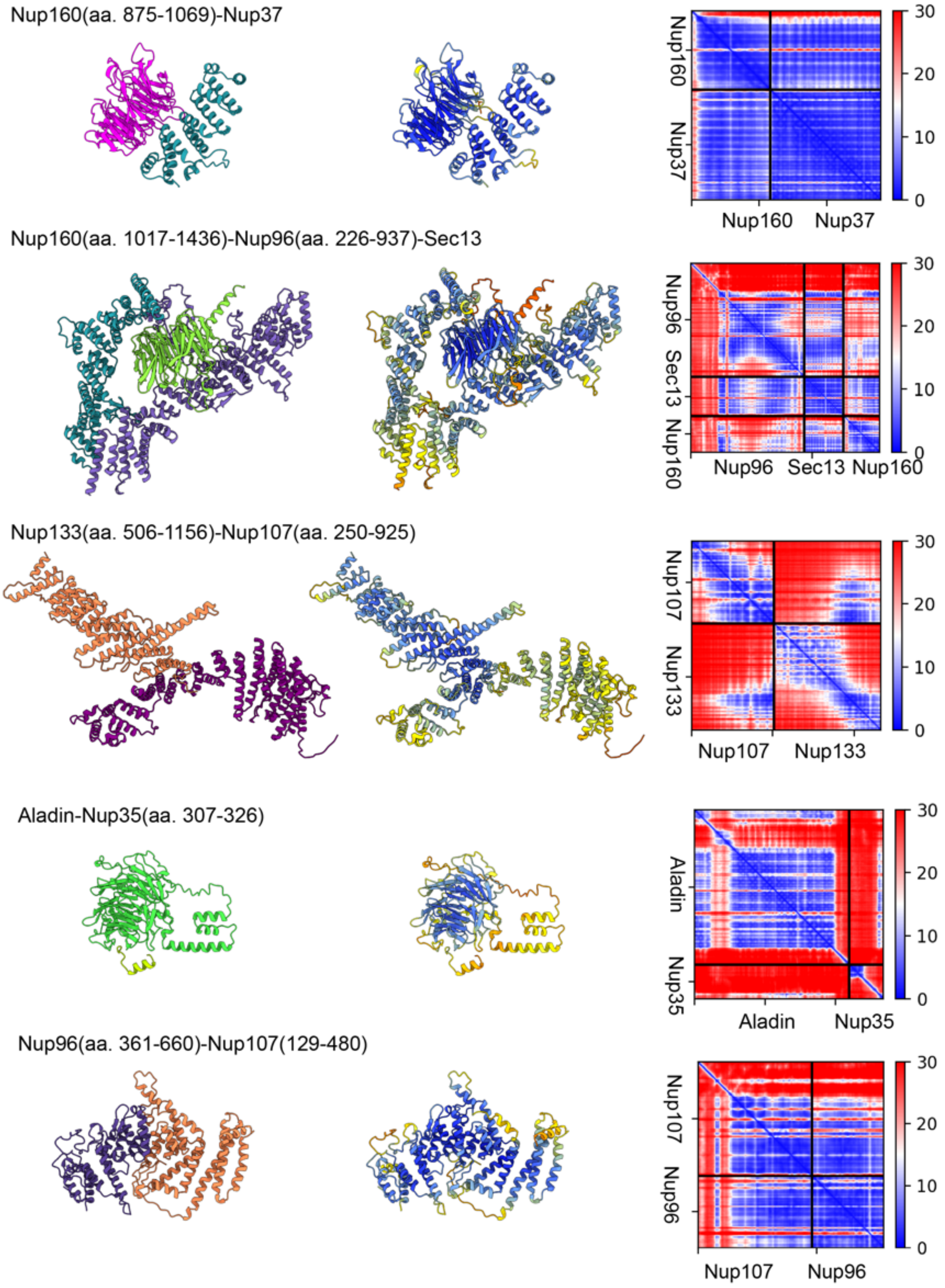
Nup160(aa. 875-1069)-Nup37
Nup160
Nup37
Nup160 Nup37
30
20
10
0
Nup160(aa. 1017-1436)-Nup96(aa. 226-937)-Sec13
Nup96 Sec13 Nup160
Nup96 Sec13 Nup160
Nup133(aa. 506-1156)-Nup107(aa. 250-925)
Nup107
Nup133
Nup107 Nup133
Aladin-Nup35(aa. 307-326)
Aladin
Nup35
Aladin Nup35
Nup96(aa. 361-660)-Nup107(129-480)
Nup107
Nup96
Nup107 Nup96

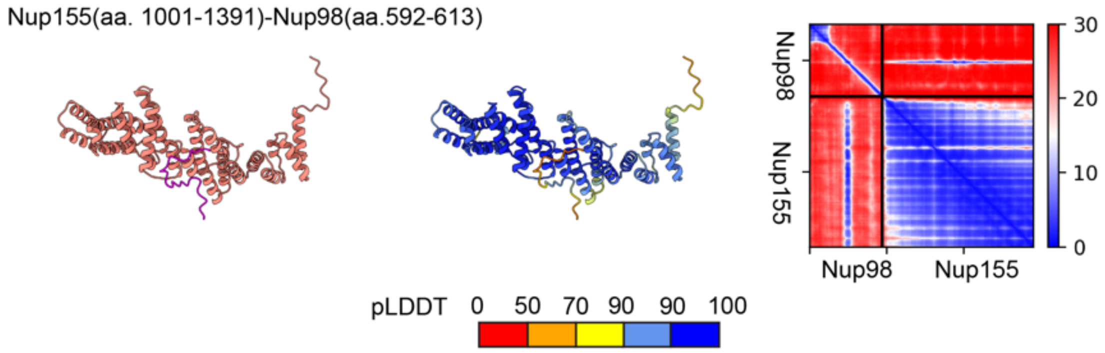


**Fig. S9 Estimated quality of structural models of human Nup sub-complexes built using AlphaFold/ColabFold**. Each row shows from left to right: (*left*) model colored by the Nup color code as in Fig. 1; (*middle*) model colored by local confidence estimated with predicted local distance difference test (pLDDT), as returned by AlphaFold. The pLDDT > 90 (dark blue) indicates high accuracy of backbone and side chain rotamers whereas pLDDT > 70 (yellow) indicates correct backbone prediction (*61*); (*right*) The confidence of inter-domain and inter-chain orientations estimated with the expected distance error in Å between all pairs of residues in the complex, as returned by ColabFold. The color at each (x, y) position of the matrix corresponds to the expected distance error in residue x's position, when the prediction and true (unknown) structure are aligned on residue y. Blue indicates low error.

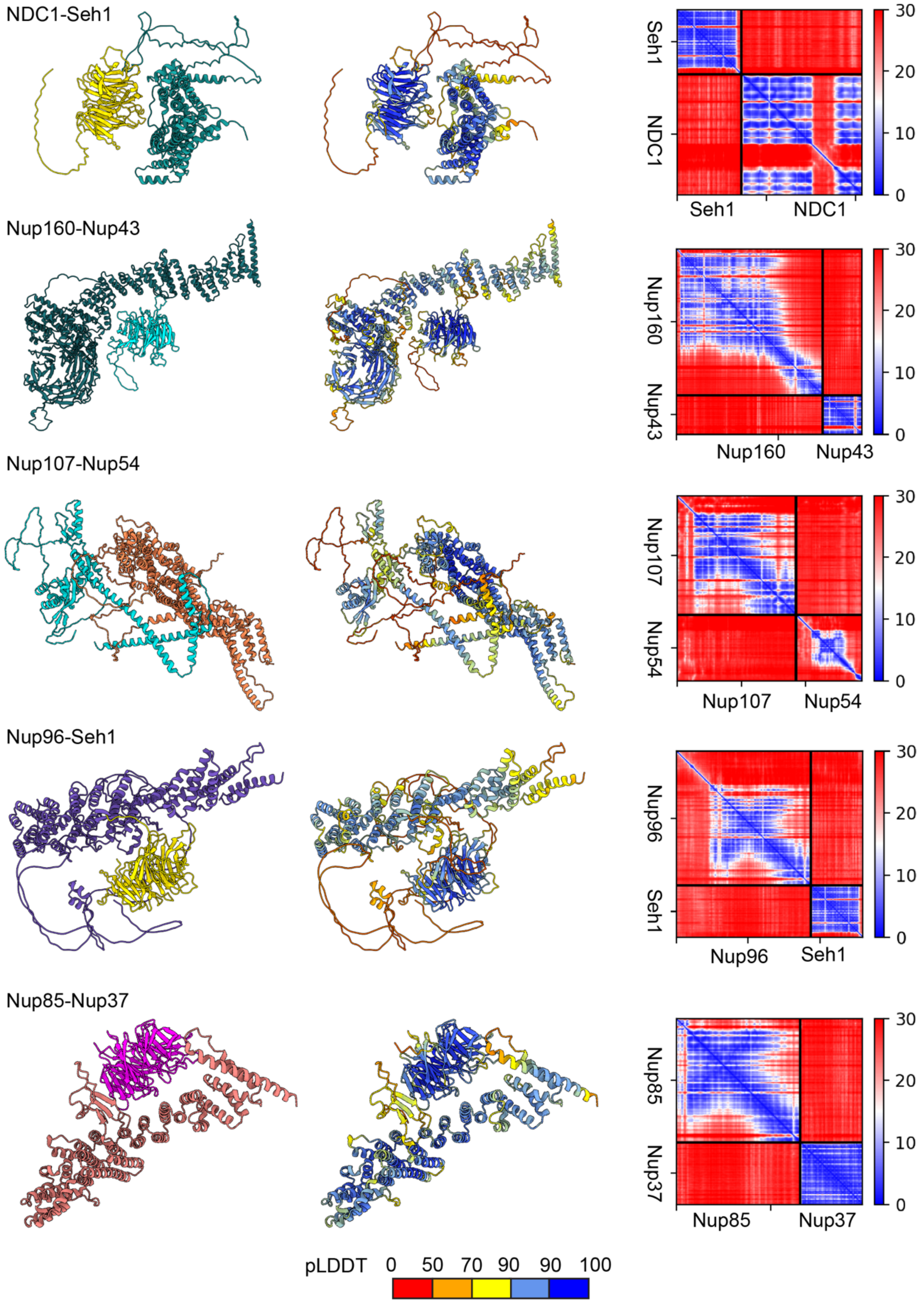


**Fig. S10 A negative control set of AlphaFold/ColabFold predictions for Nup sub-complex structures.** Various pairs of Nups not expected to interact (based on their distance in the current NPC models) were modeled. The protein chains form only loose contacts and their relative orientation is not confidently predicted as expected for true negative pairs (red areas in the plots on the right). Each row shows from left to right: (*left*) model colored by the Nup color code as in Fig. 1; (*middle*) model colored by local confidence estimated with predicted local distance difference test (pLDDT), as returned by AlphaFold. The pLDDT > 90 (dark blue) indicates high accuracy of backbone and side chain rotamers whereas pLDDT > 70 (yellow) indicates correct backbone prediction (*61*); (*right*) The confidence of inter-domain and inter-chain orientations estimated with the expected distance error in Å between all pairs of residues in the complex, as returned by ColabFold. The color at each (x, y) position of the matrix corresponds to the expected distance error in residue x's position, when the prediction and true (unknown) structure are aligned on residue y. Blue indicates low error.

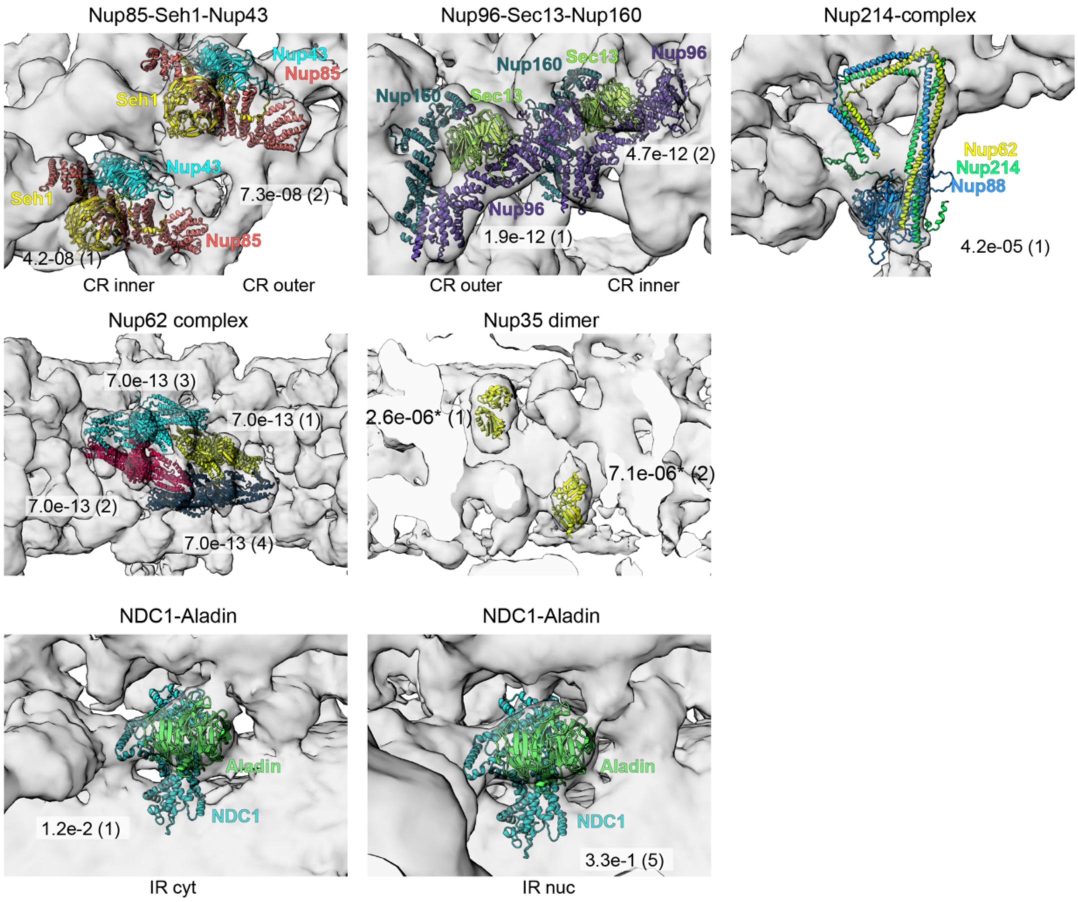


**Fig. S11 Validation of AlphaFold sub-complex models by systematic fitting to the hNPC EM maps.** Each panel shows the top fits fitted to the EM maps with proteins are colored as in Fig. 1. P-values, calculated as described in the Methods, and ranks of the fits are indicated next to the structures. All sub-complexes have been fitted to the full CR and IR maps, except NDC1-Aladin and Nup35 dimer, which were fitted to the difference map of the IR. The asterisk next to the Nup35 dimer P-values indicates uncertainty in the P-value assessment in this case due to the non-normal distribution of the fits and a long tail of P-values better than 0.05.

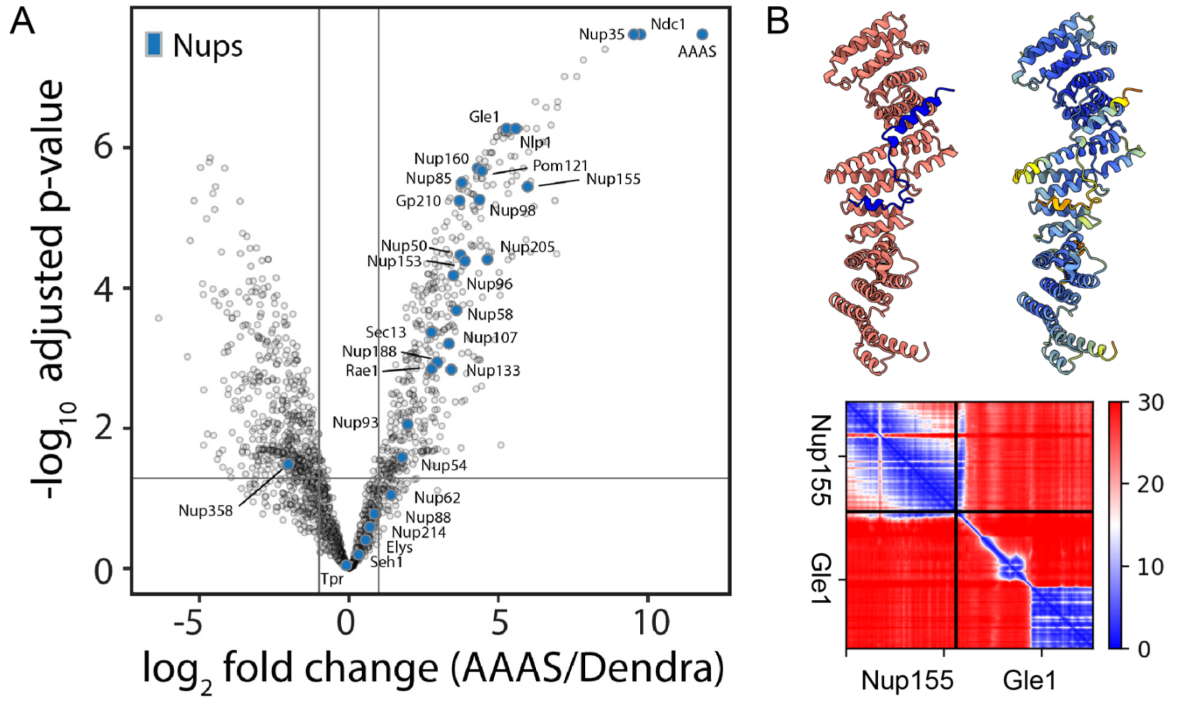


**Fig. S12 Proximity labeling experiment with BirA tagged Aladin. (A)** Volcano plot showing the enrichment of individual Nups in the proximity of Aladin (AAAS). NDC1 and Nup35 are highly enriched to similar levels as Aladin itself. Nups in proximity to the transmembrane interaction hub at the interface of the ORs with the IR are moderately enriched. This includes Nup155, GP210, Nup155, and Nup160. **(B)** The enriched nucleoporin Gle1 has been shown to interact with Nup155 (*45*), and this interaction is structurally validated by a AlphaFold/ColabFold model. In the AlphaFold model, the N terminus of Gle1 (blue) interacts with the C terminus of Nup155 (salmon, top left). The model has high-local confidence (top right) and confidence of the interaction (bottom). See Fig. S9 for the explanation of the confidence measures.

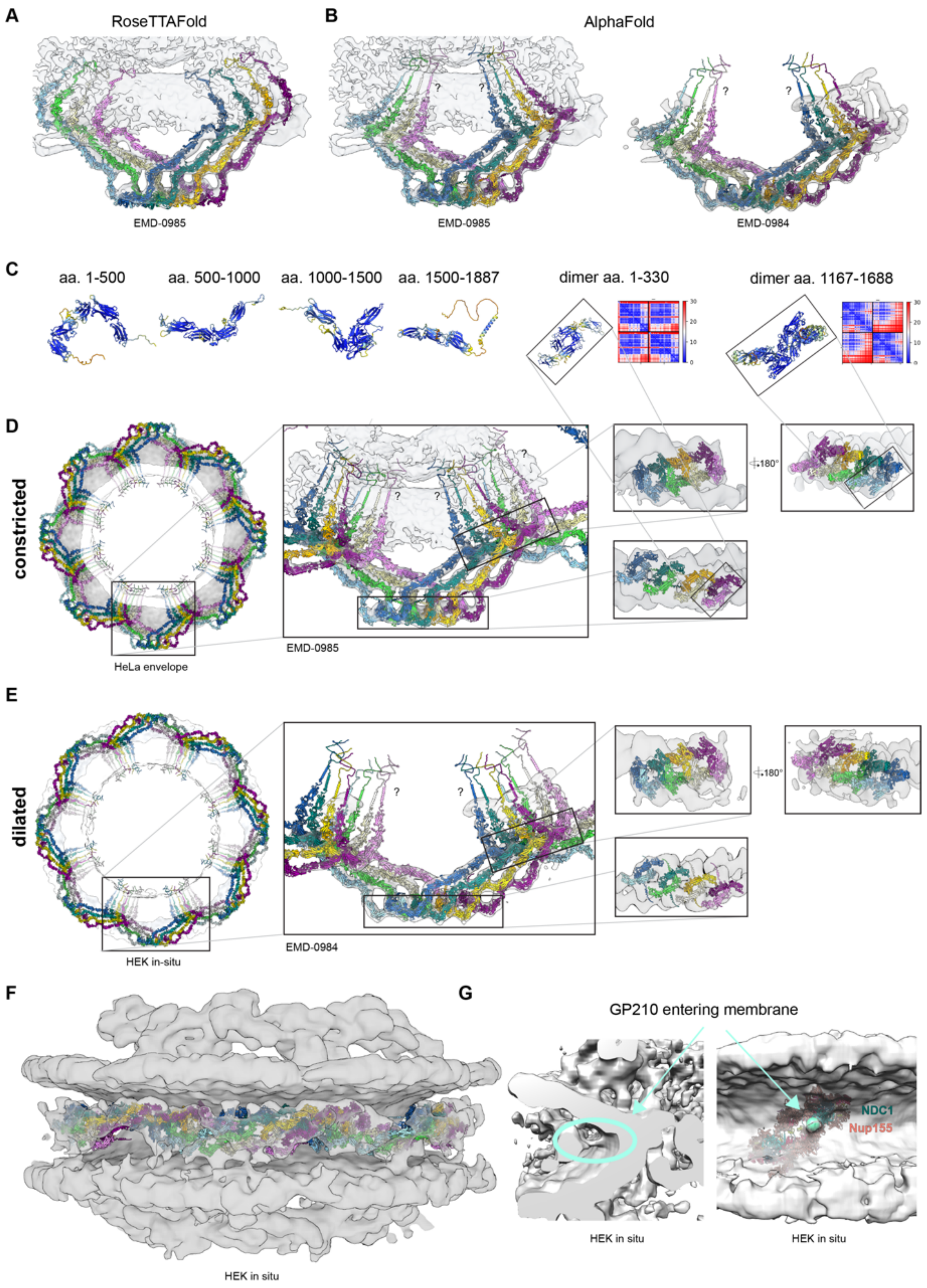


**Fig. S13 Modeling Gp210 and overview of the resulting models. (A)** A rigid-body fit of Gp210 modelled as full-length with RoseTTAfold. **(B)** AlphaFold did not produce a model fitting as a rigid body but a model could be constructed by superposing smaller AlphaFold fragments onto the RoseTTAfold fit and local refinement of the fits. **(C)** Selected fragments of AlphaFold models for monomers and dimers, which cover dimerization interfaces between neighboring copies of Gp210. The models are colored by local confidence and the heatmaps represent inter-chain accuracy as described in Fig. S9. **(D)** The full Gp210 model in the constricted NPC. Insets show various zoomed-in views at the structure. The question marks indicate that the positioning of the transmembrane helices is approximate and they could make helical bundles, which were however not predicted by AlphaFold. **(E)** The full Gp210 model in the dilated NPC. **(F)** The fit of the Gp210 model in the EM density of the dilated NPC as natively in the HEK cells. **(G)** The density connecting Gp210 density with the membrane likely encompasses the transmembrane helices of Gp210 and localizes next to the Aladin/NDC1/Nup155/Nup35 hub.

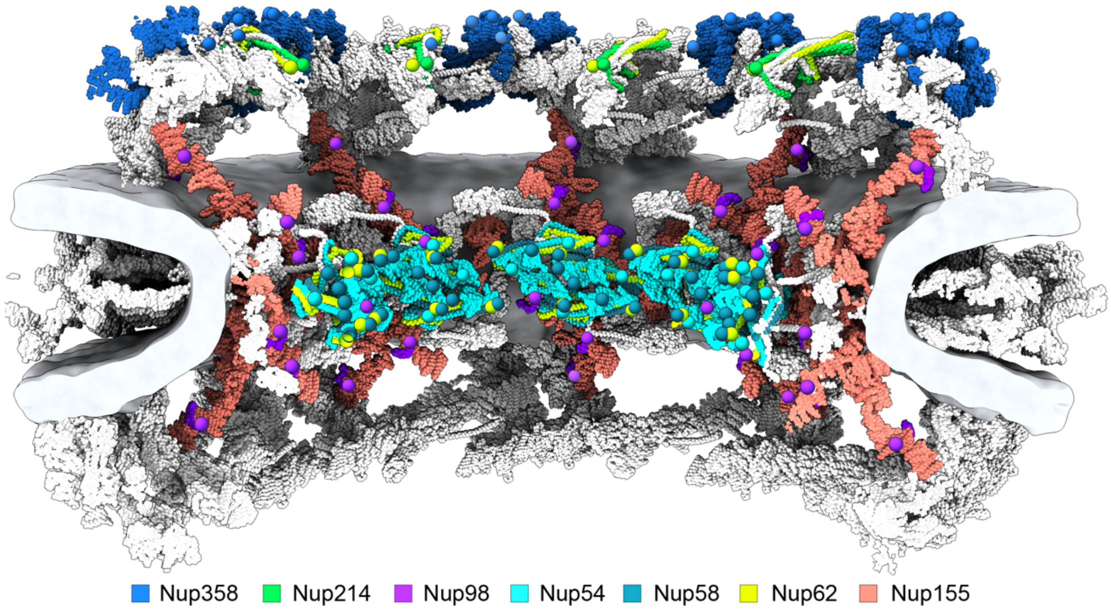


**Fig. S14 The map of FG-repeat anchoring points.** The model of the constricted NPC is shown in a cut-away view. The termini of Nup structured domains (colored) that anchor flexible FG-repeat tails (not explicitly modeled and thus not shown), are depicted as spheres with the color of the given FG-Nup. The spheres have been placed at the following residues for each Nup: Nup54 - 379, Nup58 - 246, 418, Nup62 – 332, Nup98 – 597, Nup214 – 972, Nup358 – 759.

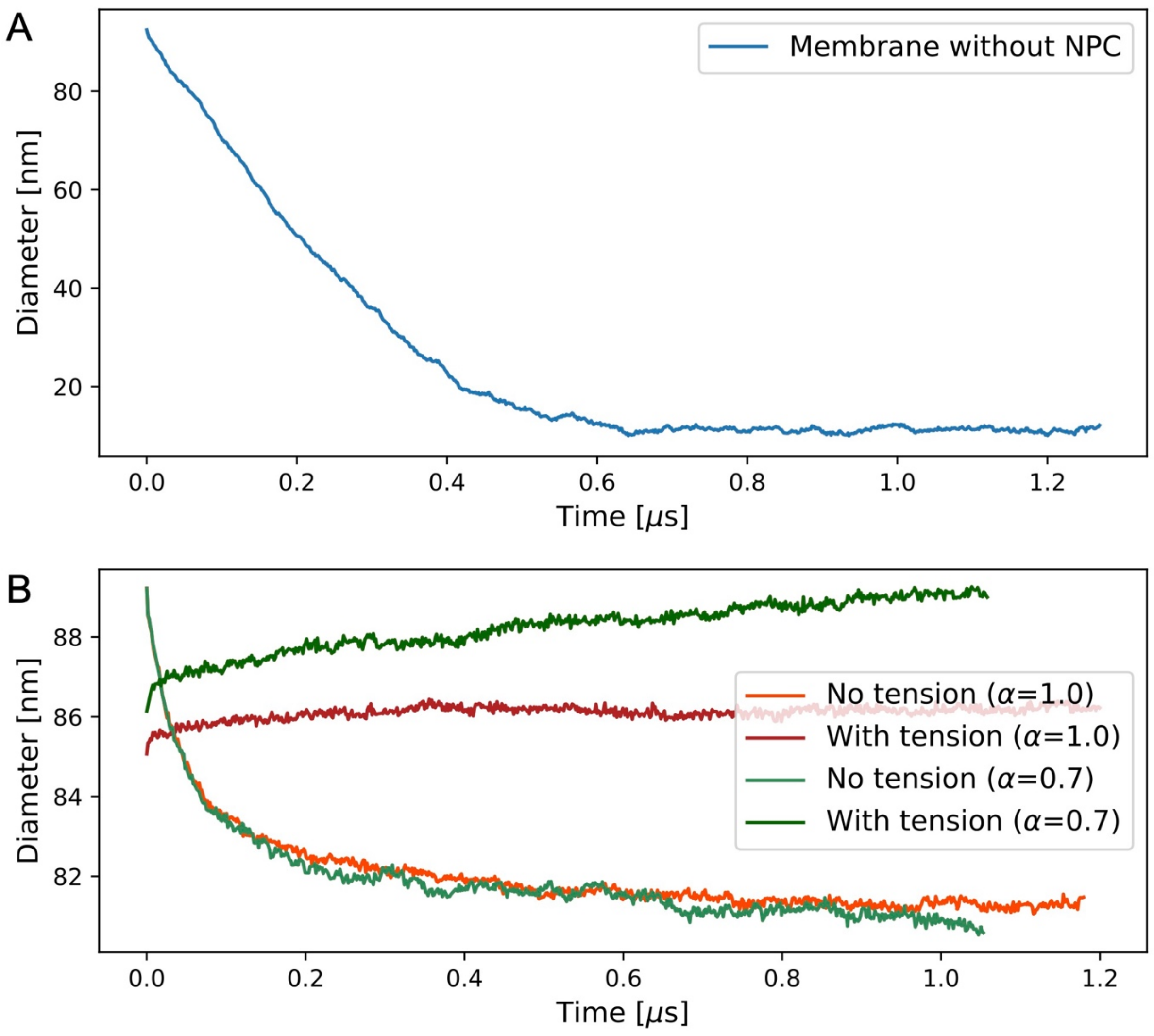


**Fig. S15** Timeseries of pore diameters. (A) Diameter of double-membrane pore in MD simulation without NPC as a function of time. See Fig. 4A for snapshots at times t=0 and 1ms. **(B)** Double-membrane pore diameter in MD simulations with hNPC scaffold. See Fig. 4B and Fig. S19 for snapshots of the simulations with $\boldsymbol{\alpha = 0.7}$ and $\boldsymbol{\alpha = 1.0}$, respectively.

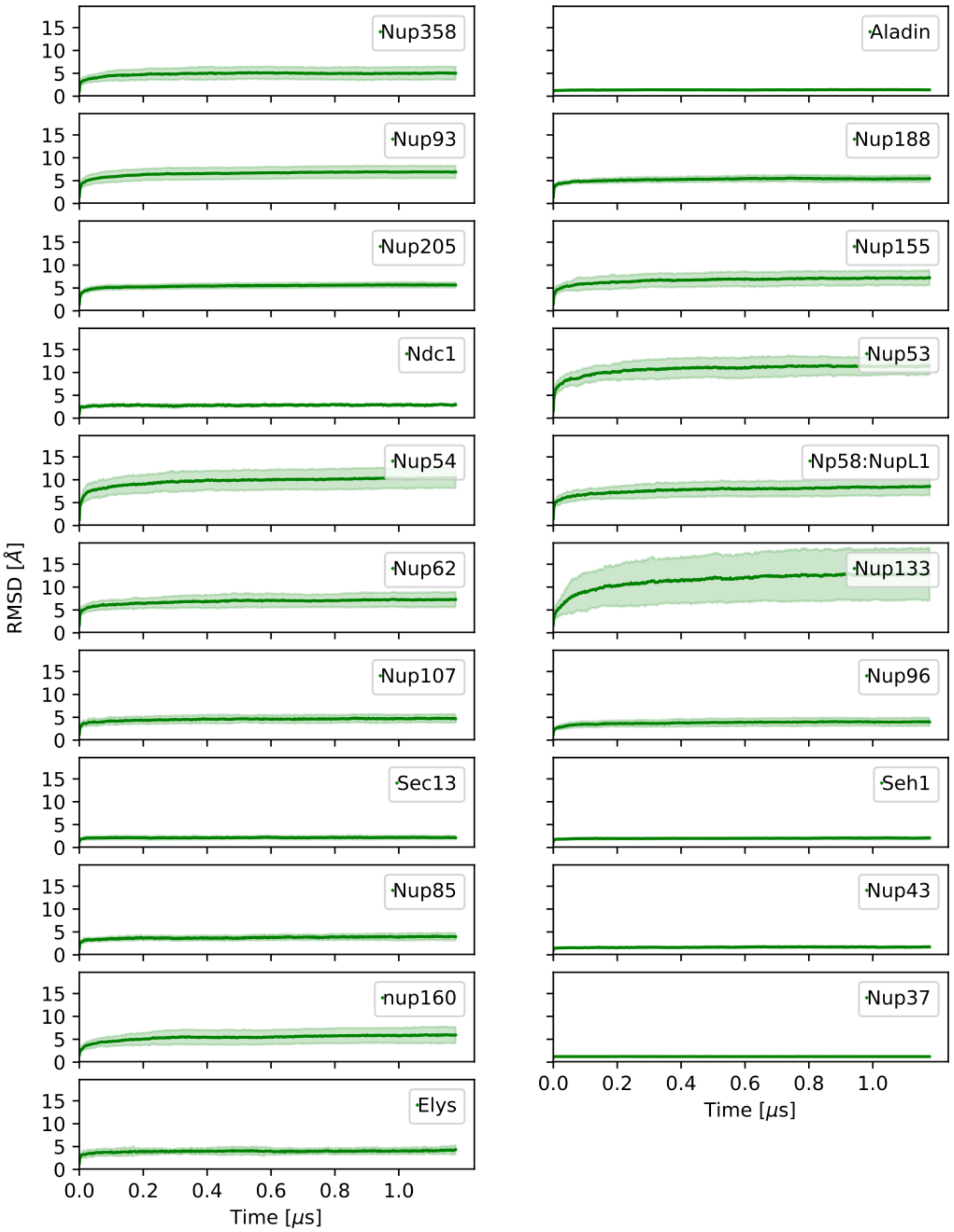


**Fig. S16 RMSD of individual proteins from their initial structures in MD simulations of the NPC at a=1.0.** The mean value of the RMSD at each given time point across all protein copies is shown as thick green line. Standard deviations are shown in shaded light green. Time points every 1.5 ns were analyzed. Protein names are indicated. The larger apparent flexibility observed in several Nups such as Nup133, Nup155, Nup35, Nup54, or Nup62 result from more extended regions or flexible linkers.

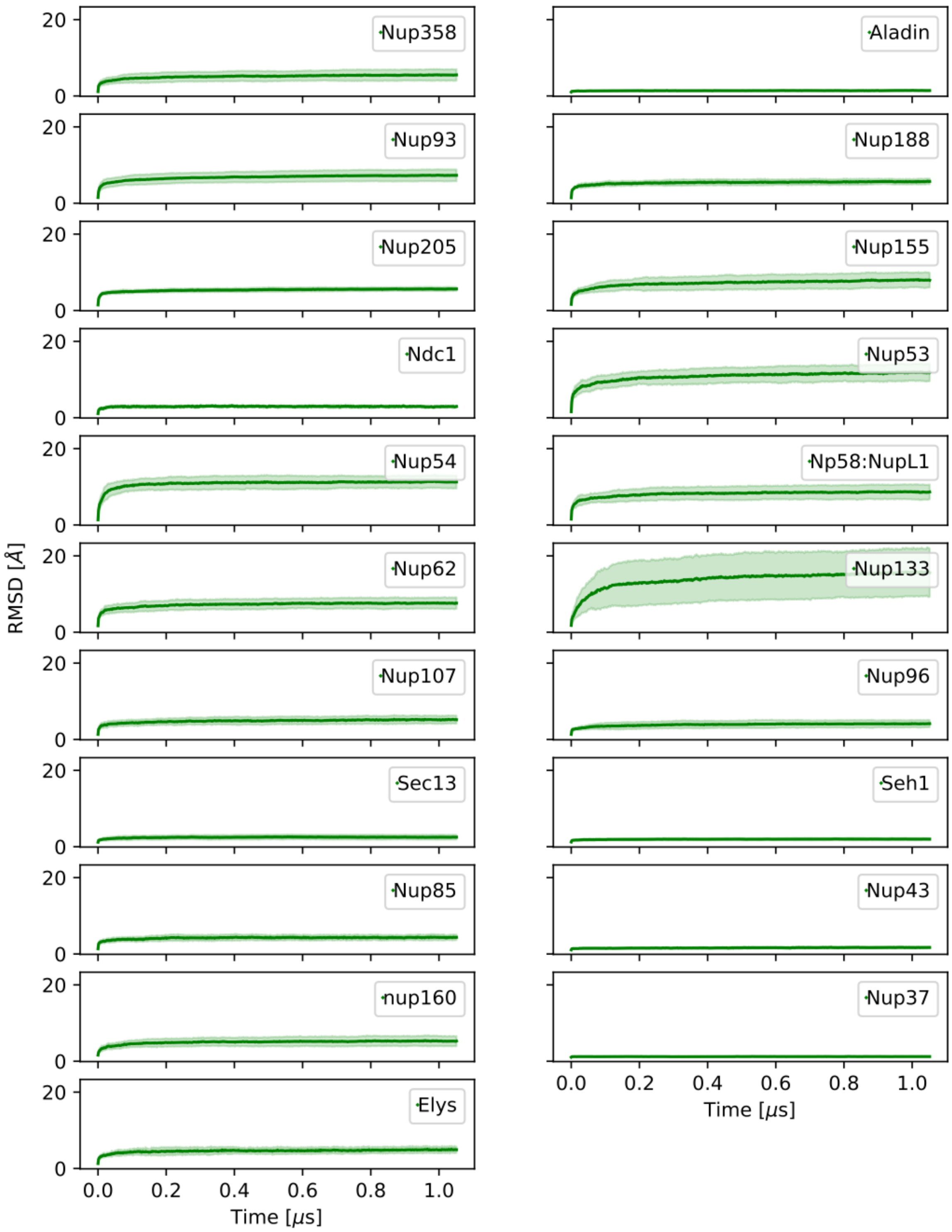


**Fig. S17 RMSD of individual proteins from their initial structures in MD simulations of the NPC at α=0.7.** The mean value at each given time point across all protein copies is shown as thick green line. Standard deviations are shown in shaded light green. Time points every 1.5 ns were analyzed. Protein names are indicated. The larger apparent flexibility observed in several Nups such as Nup133, Nup155, Nup35, Nup54, or Nup62 result from more extended regions or flexible linkers.

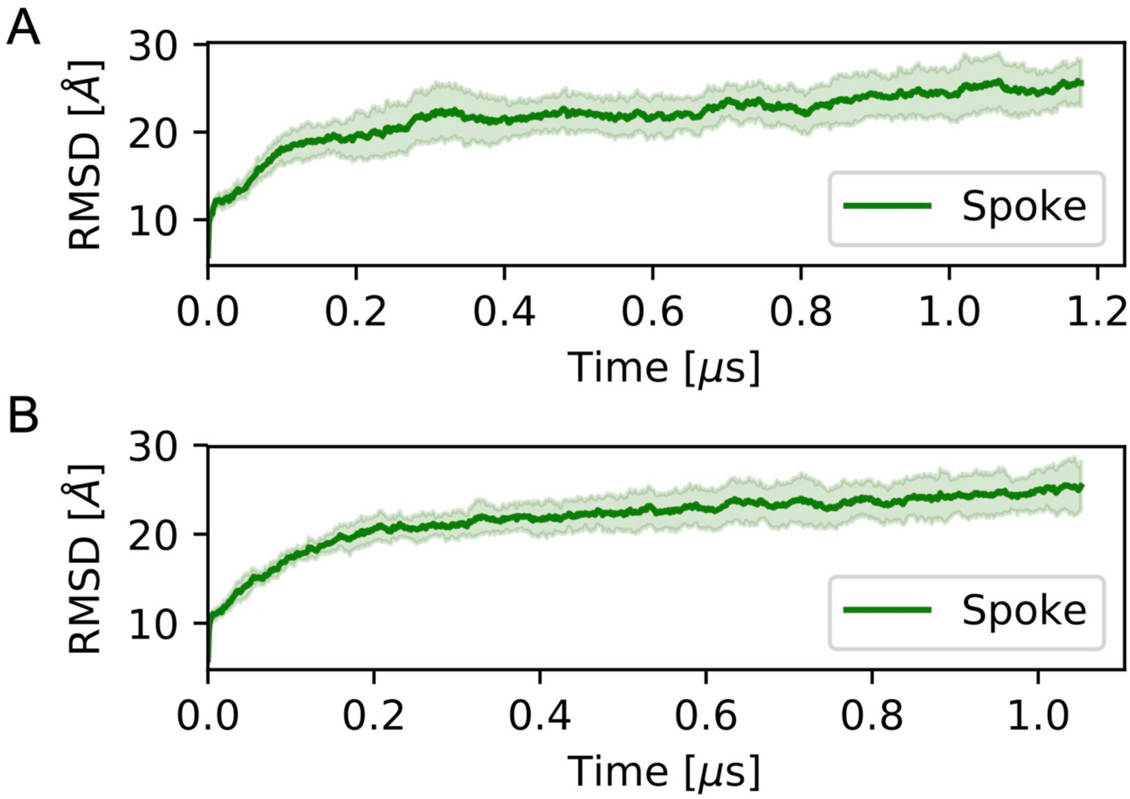


**Fig. S18 RMSD of NPC spokes from their initial structures in MD simulations of the NPC at a=1.0 and a=0.7.** The mean value of the RMSD at each given time point across all eight copies is shown as thick green line. Standard deviations are shown in shaded light green. Time points every 1.5 ns were analyzed.

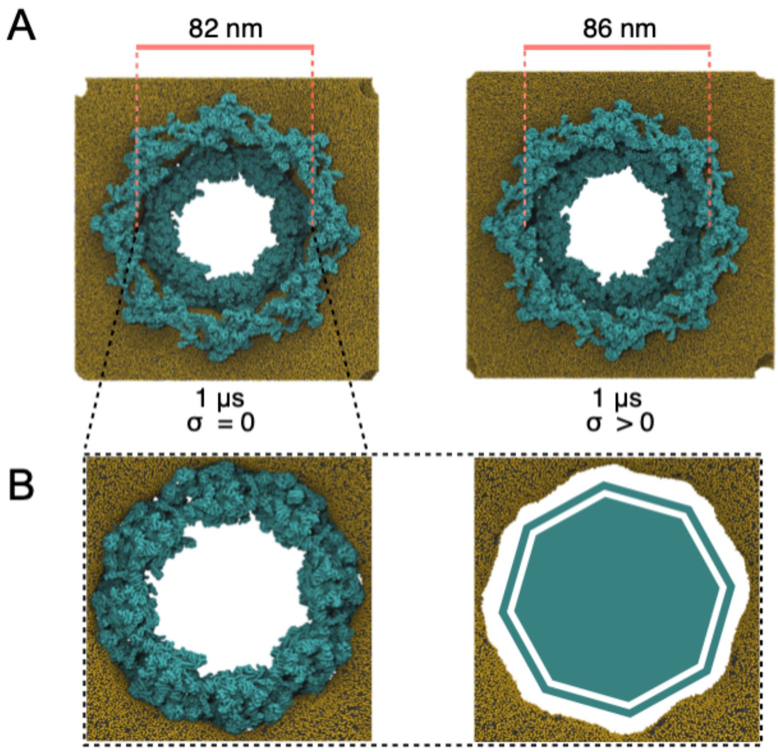


**Fig. S19 MD simulations of the complete hNPC with explicit membrane (α=1.0)**. (A) The NPC (cyan) widens by ~10% in response to lateral membrane tension (right) compared to a zero-tension simulation (left). Shown are snapshots of the relaxed structures after 1 µs of MD with unscaled protein-protein interactions (α=1). **(B)** The membrane fits tightly around the NPC inner ring (cyan, left) and forms an octagonally shaped pore (right, NPC removed).

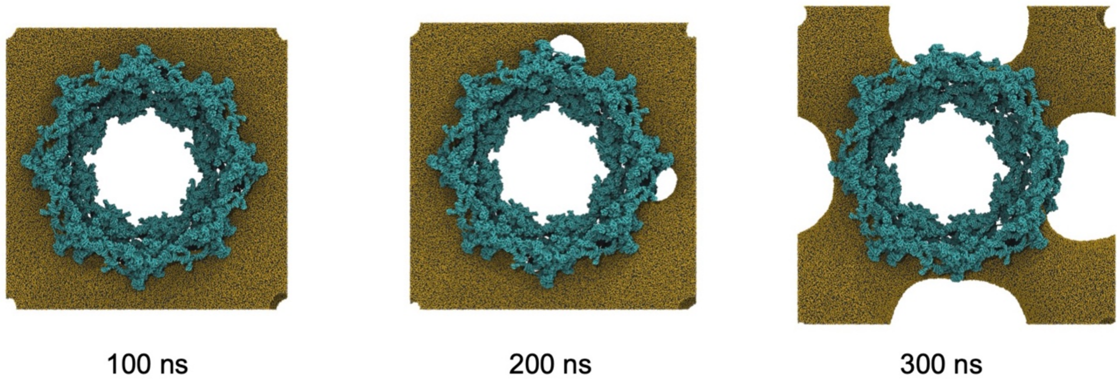


**Fig. S20 Snapshots from a representative membrane rupture event under high membrane tension ($\Delta P = 3$ bar).** The NPC scaffold backbone beads are shown in cyan. Lipid headgroups are shown as gold spheres. Water and ions are omitted for clarity.

#### Supporting Videos

**Supporting Video S1** MD simulation trajectory of an isolated half-toroidal double membrane shaped initially as in the tomographic structure of the constricted NPC. The pore tightens within 1.2 μs of MD (see also Fig. 4B and Fig. S15A for the diameter time trace). A top view of lipids is shown. Solvent is omitted for clarity.

**Supporting Video S2** MD simulation of the NPC (cyan) covering approximately 1.2 μs at $\alpha$=1.0 (see also Fig. 4B and Fig. S15A for the diameter time trace). A top view of the NPC with membrane is shown. Solvent is omitted for clarity.

**Supporting Video S3** MD simulation of the NPC (cyan) covering approximately 1.2 μs $\alpha$=0.7 (see also Fig. 4B and Fig. S15A for the diameter time trace). A top view of the NPC with membrane is shown. Solvent is omitted for clarity.
